# Mechanisms of Epigenomic and Functional Convergence Between Glucocorticoid- and IL4-Driven Macrophage Programming

**DOI:** 10.1101/2024.02.16.580560

**Authors:** Dinesh K Deochand, Marija Dacic, Michael J Bale, Andrew W Daman, Steven Z Josefowicz, David Oliver, Yurii Chinenov, Inez Rogatsky

**Author notes:** **Correspondence:** **(I.R.)**.

## Abstract

Macrophages adopt distinct phenotypes in response to environmental cues, with type-2 cytokine interleukin-4 promoting a tissue-repair homeostatic state (M2_IL4_). Glucocorticoids, widely used anti-inflammatory therapeutics, reportedly impart a similar phenotype (M2_GC_), but how such disparate pathways may functionally converge is unknown. We show using integrative functional genomics that M2_IL4_ and M2_GC_ transcriptomes share a striking overlap mirrored by a shift in chromatin landscape in both common and signal-specific gene subsets. This core homeostatic program is enacted by transcriptional effectors KLF4 and the GC receptor, whose genome-wide occupancy and actions are integrated in a stimulus-specific manner by the nuclear receptor cofactor GRIP1. Indeed, many of the M2_IL4_:M2_GC_-shared transcriptomic changes were GRIP1-dependent. Consistently, GRIP1 loss attenuated phagocytic activity of both populations *in vitro* and macrophage tissue-repair properties in the murine colitis model *in vivo*. These findings provide a mechanistic framework for homeostatic macrophage programming by distinct signals, to better inform anti-inflammatory drug design.

## Introduction

Macrophages are ubiquitous innate immune cells, named for their ability to engulf pathogens, dead cells, cell debris, and other foreign objects and initiate inflammatory and immune responses ^11^. More recently, macrophages were recognized to broadly contribute to the maintenance of homeostasis through numerous, often tissue-specific functions, including phagocytosis of apoptotic cells in diverse tissues, iron recycling in the spleen and liver, and erythropoiesis in the bone marrow and fetal liver ^2^. Macrophages differ ontogenically and principally originate from three sources during embryonic and fetal development: the yolk sac, the fetal liver and the bone marrow ^3^. In addition to developmental imprinting, macrophages are extremely plastic, with various signals in their microenvironment ^4^, such as cytokines and growth factors, shaping distinct populations ^4, 5, 6^. The ability of macrophages to adopt specialized phenotypes in response to different stimuli has been termed polarization ^7^ and is central to many physiological and pathological processes including wound healing, metabolic abnormalities and cancer ^8^.

A seminal study of macrophage gene expression patterns upon exposure to a diverse range of stimuli in vitro revealed that their transcriptomes exist along a spectrum of states; the opposite poles were represented by those enacted by proinflammatory lipopolysaccharide (LPS) and interferon gamma (IFNγ) vs. type-2 cytokines IL4, 9 and 13, corresponding to historically described M1-(inflammatory) and M2-like (homeostatic) states, respectively ^9^. One of the most well-studied pathways leading to the M2 phenotype is triggered by the cytokine IL4 (hereafter, M2_IL4_) ^10, 11^. IL4 binds to the receptor subunit IL4Rα ^12^, leading to the IL4 receptor complex assembly and phosphorylation, followed by phosphorylation and nuclear translocation of the transcription factor STAT6 and induction of many STAT6 targets ^12, 13^. These include transcription factors EGR2 and PPARγ, as well as effector molecules: Chil3, involved in nematode clearance, Arg1, necessary for arginine to ornithine conversion thus supporting collagen production and repair, and Mrc1, needed for phagocytosis of pathogens ^14^. Kruppel-like factor 4 (KLF4), another transcriptional regulator and a downstream target of STAT6, is a key player in M2 polarization, critical for the upregulation M2 and downregulation of inflammatory M1 genes ^15^. Consistently, myeloid cell-specific KLF4 deletion in mice led to metabolic inflammation, predisposing them to diet-induced obesity, glucose intolerance, and insulin resistance, as well as impaired wound healing *in vivo* ^15^. More recently, it has been shown that EGR2 bridges the early and late gene expression programs during M2 polarization ^14^.

A distinct stimulus that promotes the homeostatic phenotype in macrophages are the glucocorticoids (GCs) – steroid hormones produced by the adrenal cortex that have remained the cornerstone of anti-inflammatory therapies for nearly 70 years ^16, 17, 18, 19^. GCs activate the glucocorticoid receptor (GR), a ligand-dependent transcription factor of the nuclear receptor superfamily which, among hundreds of targets, represses pro-inflammatory and activates anti-inflammatory genes ^20, 21^. GR functions by engaging various coregulators including GR-interacting protein 1 (GRIP1, NCoA2) which participates in both processes ^22, 23, 24, 25^. With respect to GC-polarized M2 (hereafter M2_GC_), glucocorticoid-induced leucine zipper (GILZ, *Tsc22d3*), a well-known GR target, enhances efferocytosis by macrophages, a hallmark of M2 activity, both *in vitro* and *in vivo* ^26^. Several other genes – *CD206, CD163, MerTK* – known to be highly expressed in M2 – are reported GR transcriptional targets ^27, 28, 29^. Nonetheless, the precise roles of GR or GRIP1 in generating the M2_GC_ state have not been studied. Interestingly, KLF4 is itself a GR-regulated gene in macrophages ^30^, suggesting that the M2_IL4_ and M2_GC_ macrophage states are inherently connected. However, the interplay between KLF4 and GR in establishing the homeostatic state remains unexplored.

Here, we evaluate the transcriptomic and epigenomic landscape of the M2_GC_ and M2_IL4_ populations of primary mouse macrophages. We comprehensively assess the extent of the functional convergence vs. distinctions between these differentially polarized macrophages at the level of transcription factor binding, chromatin state and gene expression. Finally, we establish a shared transcriptional cofactor utilization as the underlying mechanism in the programming of homeostatic macrophages *ex vivo* and *in vivo*, in the mouse model of tissue damage.

## Results

### Overlapping yet distinct transcriptomic profiles of the M2_IL4_ and M2_GC_ macrophage populations

To evaluate the transcriptomic make-up of M2_IL4_ and M2_GC_ macrophages, we performed RNAseq on mouse primary bone marrow-derived macrophages (BMDM) exposed to IL4 or GCs: corticosterone (Cort, endogenous GC in rodents) and dexamethasone (Dex, highly potent, synthetic, clinically used GC) for 24 h. Transcriptome analysis revealed 720, 154, and 307 differentially expressed genes (DEGs) in M2_IL4_, M2_Cort_, and M2_Dex_, respectively, relative to M0 (fold change (FC) ≥ 2; FDR <0.05) (Fig. 1a). As expected, the M2_Cort_ transcriptome was fully included in that of M2_Dex_, and even though 157 of Dex-regulated genes failed to pass the significance threshold in M2_Cort_ at n=3, they displayed a similar pattern of regulation, as shown for representative genes (Extended Data Fig. 1a). This is consistent with Dex being a more stable and potent GR ligand than Cort. Importantly, whereas 628 and 215 genes were specific to M2_IL4_ and M2_Dex_ macrophages, respectively, 92 genes – 30% of all DEGs in the M2_Dex_ population – were also differentially expressed and changing in the same direction in the M2_IL4_.

**Fig. 1.**
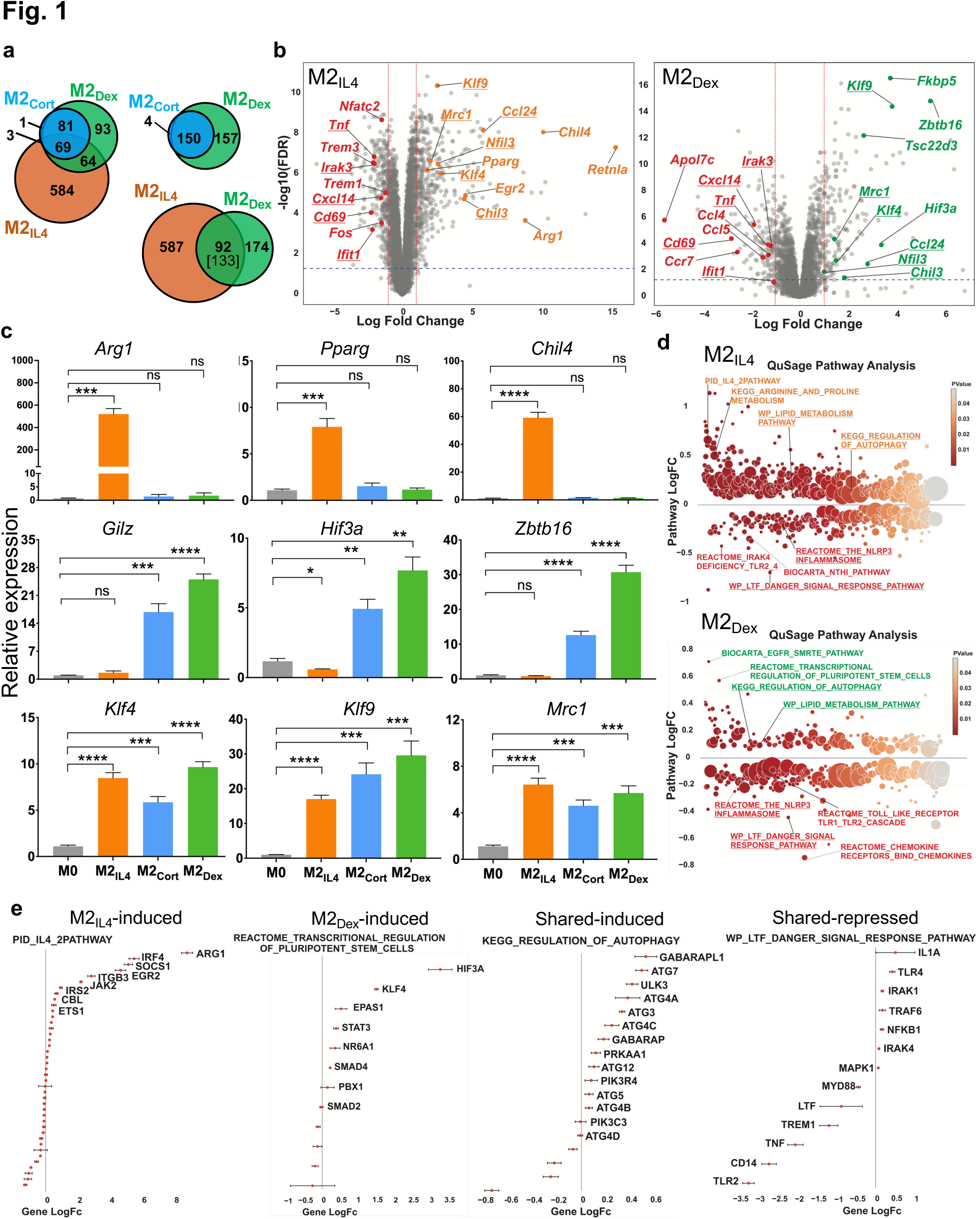
| Transcriptomic analysis of the M2_IL4_ and M2_GC_ populations. **a**, Transcriptomic changes in M0 macrophages polarized with IL4 or GCs corticosterone (Cort) or dexamethasone (Dex) for 24 h were determined by RNAseq (n=3). Venn diagrams show DEGs in M2_IL4_ and M2_GC_ relative to M0 (FC ≥ 2; FDR <0.05); 92 of 133 genes are regulated in both M2_IL4_ and M2_Dex_ in the same direction. **b,** Volcano plots show DEGs in M2_IL4_ (left) and M2_Dex_ (right) relative to M0 (LogFC ≥ 1; FDR< 0.05). Selected upregulated genes are highlighted in orange (M2_IL4_) and green (M2_Dex_). Downregulated genes are highlighted in red in both populations. Examples of the M2_IL4_:M2_Dex_ shared DEGs are underlined. **c,** RT-qPCR validation of genes upregulated selectively in M2_IL4_ (top), M2_GC_ (middle), or both (bottom). Student’s t-test; *p < 0.05, **p < 0.01, ***p < 0.001, ****p < 0.0001, ns, non-significant. n=4, error bars are SEM. **d,** Differentially regulated pathways (unadjusted p < 0.01) identified using QuSAGE and MsigDB canonical pathways (c2.cp.v7.3, Broad Institute) are shown for M2_IL4_ (top) and M2_Dex_ (bottom). Underlined are the shared pathways between M2_IL4_ (upregulated, orange; downregulated, red) or M2_Dex_ (upregulated, green; downregulated, red). Circle size is proportional to the number of genes in the pathway and color signifies p-value. **e,** Genes from indicated pathways induced or repressed in M2_IL4_, M2_Dex_, or both are plotted LogFC±SD and ordered by p-value.

Among DEGs upregulated in M2_IL4_ (n=431) and M2_Dex_ (n=161), we found shared genes characteristic of the M2-like phenotype, encoding proteins with the anti-inflammatory, tissue homeostasis and repair functions (Fig. 1b, underlined: *Klf4*, *Klf9*, *Chil3*, *Ccl24*, and *Mrc1*). However, in each population, we also noted genes that were more signal-specific: *Arg1*, *Chil4*, *Pparg*, *Egr2*, and *Retnla* for M2_IL4_ and *Fkbp5*, *Hif3a*, *Tsc22d3*, and *Zbtb16* for M2_Dex_ (Fig. 1b and Extended Data Fig. 1b). The M2_IL4_-specific targets are well-established markers of the M2_IL4_ state; the M2_Dex_-specific genes include canonical GR targets. We validated genes induced selectively in M2_IL4_ (*Arg1*, *Pparg*, and *Chil4*), M2_GC_ (*Tsc22d3 (Gilz)*, *Hif3a*, and *Zbtb16*) or both (*Klf4*, *Klf9*, and *Mrc1*) by quantitative PCR (Fig. 1c). M2 polarization often involves the suppression of the pro-inflammatory arm of gene expression. Indeed, the pool of downregulated genes (289 in M2_IL4_ and 146 in M2_Dex_) included components of the type 1 interferon network, chemokines, and pro-inflammatory mediators (*Tnf, Cxcl14, Ifit1* and *Irak3*, common for M2_IL4_ and M2_GC_, and *Ccl4* and *Ccl5,* specific for M2_GC)_ typically upregulated in inflammatory macrophages (Fig. 1B and Extended Data Fig. 1b).

Concomitant induction of shared homeostatic genes and dampened expression of M1-specific ones suggests that both IL4– and Dex-mediated macrophage polarization may rely on overlapping set of regulatory pathways. Quantitative Set Analysis of Gene Expression (QuSage) revealed 677 and 364 pathways regulated in M2_IL4_ and M2_Dex_ populations relative to M0 (p<0.05; Extended Data Fig. 1c; Extended Data Table 1). Of those, 502 and 189, respectively, were M2_IL4_– and M2_Dex_-specific, whereas 142 pathways differentially regulated in M2_Dex_ were also regulated in M2_IL4_ and in the same direction. Upon stratifying the shared pathways by the direction of change, 49 pathways were upregulated in both M2_IL4_ and M2_Dex_, and 93 pathways were downregulated in both (Extended Data Fig. 1c), indicating that overlapping set of pathways were engaged during polarization by IL4 or GCs. Shared upregulated pathways in M2_IL4_ and M2_GC_ included Lipid Metabolism, Regulation of Autophagy, Negative Regulation of FLT3 and MET activities and MTOR signaling (Fig. 1d, Extended Data Fig. 1d, underlined; Extended Data Table 1). Among signal-specific pathways, the IL4 and Arginine and Proline metabolism-related pathways were selectively upregulated in M2_IL4_, whereas EGFR_SMRTE pathway and Transcriptional Regulation of Pluripotent Stem Cells were specific to M2_Dex_ and M2_Cort_ (Fig. 1d, Extended Data Fig. 1d and Extended Data Table 1). As expected, multiple inflammatory pathways including the NLRP3 inflammasome, CD209 signaling, Toll-like receptor (TLR) cascades, FLMP and LTF danger signal response pathways were downregulated in both M2_IL4_ and M2_GC_ (Fig. 1d-e and Extended Data Fig. 1d), consistent with the immuno-suppressive and anti-inflammatory phenotypes of M2 macrophages (or the effect of IL4 or GC treatment).

Previously validated signal-specific genes were found in M2-specific pathways. For example, *Arg1* and *Egr2* were upregulated specifically in M2_IL4_, whereas genes involved in transcriptional regulation of pluripotent stem cells such as *Hif3a* and *Klf4* were upregulated in the M2_Dex_, confirming that the key population-specific markers are associated with functionally relevant pathways (Fig. 1e). Notably, downregulated pathways that were signal-specific, such as the NFkB-related NTHI pathway in the M2_IL4_ (encompassing *Tnf*, *Nfkbia*, *Myd88*, and *RelA*) or Chemokine and Chemokine receptors in M2_GC_ population (including *Ccl3*, *Ccl4*, *Ccl5*, *Cx3cr1*, *Ccr7*, *Cxcl2*) (Extended Data Fig. 1e) were still functionally related to inflammation and innate immune responses. Collectively, these findings indicate the existence of overlapping gene expression programs elicited in the M2_IL4_ and M2_GC_ broadly related to anti-inflammatory, homeostatic and tissue reparative functions.

### IL4 and GCs drive overlapping changes in the macrophage enhancer landscape

To investigate the mechanism underlying transcriptomic overlap between M2_IL4_ and M2_GC_ macrophage populations, we surveyed accessible chromatin regions using ATACseq. The genomic distribution of ATACseq peaks was comparable in M0, M2_IL4,_ and M2_Dex_ with accessible chromatin mainly observed in the gene body (introns and exons), intergenic regions and promoters (Extended Data Fig. 2a). Out of 71,942 peaks total, we detected 31,654 and 16,379 differential peaks in the M2_IL4_ and M2_Dex_ populations, respectively, relative to M0 (FC ≥ 2; FDR =0.05; Fig. 2a). A two-fold higher number of the differential sites in M2_IL4_ is as expected based on the larger transcriptomic changes occurring in response to IL4 compared to GCs. Out of all differential sites, 23,441 and 8,166, respectively, were M2_IL4_– and M2_Dex_-specific (Fig. 2a) with a relatively even distribution between up– and downregulated peaks (Fig. 2b). Interestingly, 4,913 of differentially regulated peaks in the M2_Dex_ population were also regulated in M2_IL4_ and in the same direction (Fig. 2a), consistent with an observed overlap in the transcriptomic profiles of the M2_IL4_ and M2_GC_ populations.

**Fig. 2.**
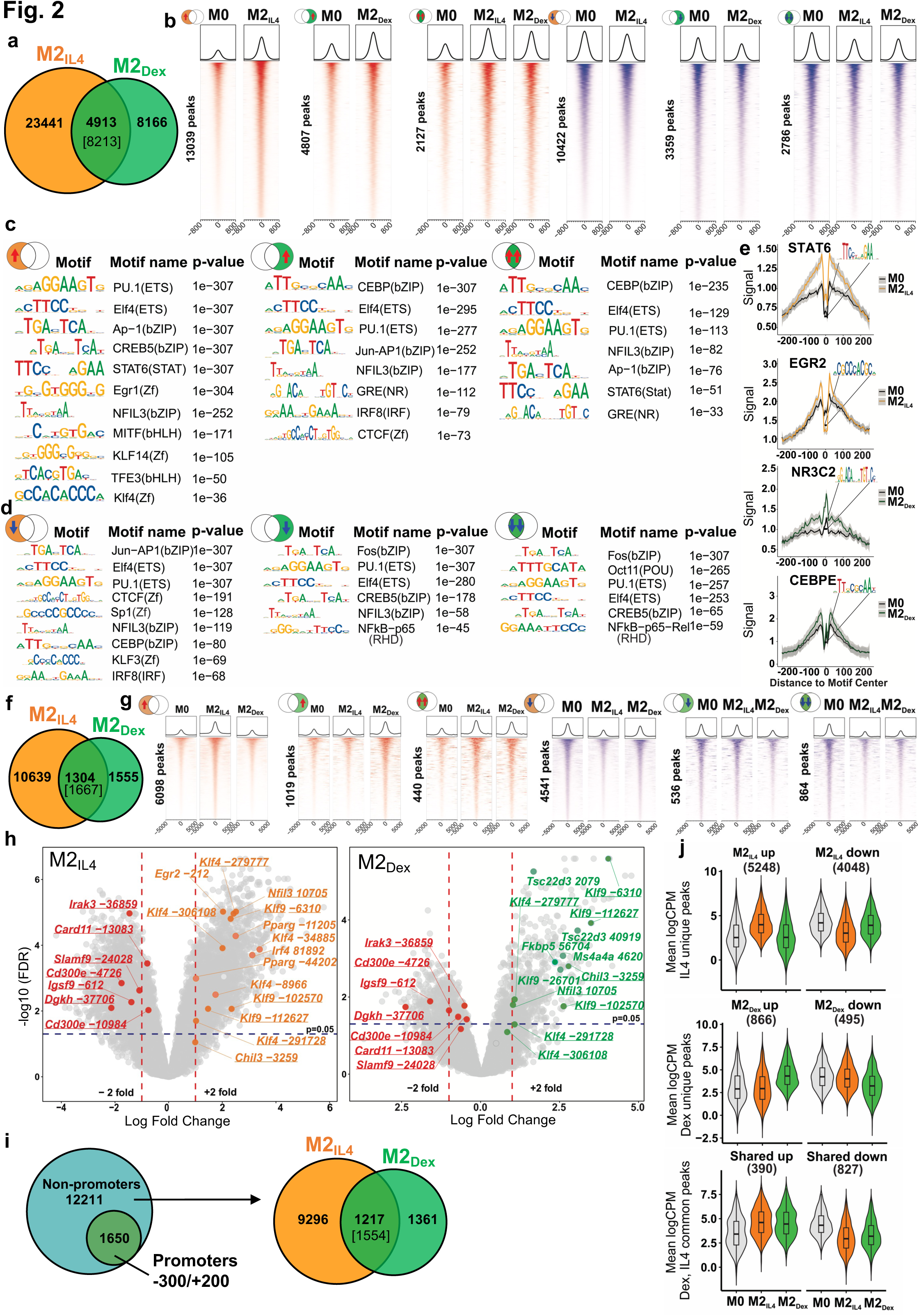
| M2IL4 and M2_GC_ macrophage populations share epigenetic landscape. **a**, ATACseq was performed on M0, M2_IL4_ and M2_Dex_. Venn diagram shows the number of differential ATACseq peaks in M2_IL4_ and M2_Dex_ relative to M0 (n=4, FC ≥ 2; FDR <0.05). 4,913 of 8,213 peaks were differential in both populations and regulated in the same direction. **b,** Heatmaps of the induced (red) and suppressed (blue) signal-specific and shared M2_IL4_ and M2_Dex_ ATACseq peaks (FC ≥ 2; FDR <0.05). **c-d,** Enriched transcription factor binding motifs with associated p-values were identified by HOMER known motif analysis with JASPAR 2022 motif database in signal-specific and shared up-(**c**) and downregulated (**d**) ATACseq peaks in M2_IL4_ and M2_Dex_. **e,** Polarization-induced increases in the 5’ Tn5 cut site counts within indicated binding motifs of interest in the M2_IL4_ and M2_Dex_ populations. **f,** Venn diagram shows the differential M2_IL4_ and M2_Dex_ H3K27ac ChIPseq peaks relative to those in M0 (FDR< 0.05) with 1,304 of 1,667 peaks regulated in the same direction in both populations. **g,** Heatmaps of induced (red) and suppressed (blue) signal-specific and shared filtered H3K27ac peaks (FC ≥ 2; FDR <0.05) in M2_IL4_ and M2_Dex_. **h,** Volcano plots show differential H3K27ac ChIPseq peaks for M2_IL4_ (left) and M2_Dex_ (right) relative to those in M0 (LogFC ≥ 1; FDR< 0.05); the location of selected hyperacetylated sites relative to the TSS of the closest gene are shown in orange (M2_IL4_) or green (M2_Dex_). Hypoacetylated sites are shown in red for each population. Sites overlapping in M2_IL4_ and M2_Dex_ are underlined. **i,** Non-promoter 12,211 H3K27ac peaks after excluding 1,650 promoter-proximal peaks (−300/+200 relative to TSS) are shown as a Venn diagram with the number of differential, relative to M0, M2 population-specific and shared peaks (FC ≥ 2; FDR<0.05) indicated. **j,** Average H3K27ac ChIPseq signals at M2_IL4_ (top), M2_Dex_ (middle), and shared (bottom) differential non-promoter peaks from (I) represented as violin plots.

Specifically, we identified 13,039 and 4,807 chromatin regions that became more accessible only in M2_IL4_ or M2_Dex_, respectively, and 2,127 peaks that ‘opened’ in both M2-like populations (Fig. 2b, red). At the same time, 10,442 peaks in M2_IL4_, 3,359 peaks in M2_Dex_, and 2,786 peaks in both populations (blue) lost chromatin accessibility. These findings indicate that IL4 and GC impart broad changes in chromatin during polarization, with a subset of sites affected in both populations. Transcription factor binding motif enrichment analysis was performed for the ATACseq peak subsets separated by the direction of change and stimulus. A group of motifs from the AP1, ETS and C/EBP families were enriched in all peak subsets (Fig. 2c-d) likely representing lineage-determining transcription factors involved in cell proliferation and differentiation ^31^. Binding motifs for transcription factors that orchestrate the M2 polarization in response to IL4, namely STAT6, KLF4 and EGR1/2, were enriched in M2_IL4_ accessible peaks (Fig. 2c). Similarly, the GR-specific GRE motif is enriched, as expected, in the M2_Dex_ peak set with increased accessibility. In shared upregulated peaks, both STAT6 and GRE motifs were enriched. Furthermore, a binding motif for NFIL3, a transcription factor upregulated by both IL4 and GC signaling, was enriched in all 3 peak subsets with increased accessibility in polarized macrophages. Importantly, analysis of Tn5 cut site distribution near overrepresented motifs (Fig. 2e) revealed changes in site accessibility in both M2_IL4_ (STAT6, EGR2) and M2_Dex_ (NR3C2/GRE, CEBPE) indicating that the observed enrichment was functional. Interestingly, NFkB-p65 motif appeared in M2_Dex_ and M2_IL4_:M2_Dex_ shared downregulated clusters (Fig. 2d), consistent with the well-established repression of the NFkB activity by GR ^21^. Together, motif enrichment analysis suggested enhanced binding of M2-specific transcription factors in opening chromatin sites, while motifs for inflammation-driving factors were found in less accessible chromatin in both signal-specific and shared peak subsets.

Histone modifications are typically more stable than transcription factor binding and indicate longer-lived activating or repressive changes in chromatin state ^32^. Among them, acetylation of histone H3 lysine 27 (H3K27ac) is commonly recognized as an enhancer mark found in active chromatin regions ^33^. Thus, we performed chromatin immunoprecipitation (ChIPseq) for H3K27ac in M0, M2_IL4_ and M2_GC_ to investigate polarization-induced acetylation changes at enhancers and/or promoters. Differential analysis of H3K27ac data shows that 12,298, 1,192, and 3,222 peaks were regulated in M2_IL4_, M2_Cort_, and M2_Dex_ populations, respectively, relative to M0 (Extended Data Fig. 2b). As expected, nearly all differential peaks detected in M2_Cort_ overlapped with those in M2_Dex_, consistent with transcriptomic changes, and we therefore used only M2_Dex_ for subsequent analyses. Polarization with IL4 and Dex produced 10,639 and 1,555, respectively, treatment-specific differential H3K27ac peaks (Fig. 2f). In addition, we identified 1,667 peaks overlapping in M2_IL4_ and M2_Dex_ populations, of which 1,304 (40% of total M2_Dex_ differential peaks) were regulated in the same direction (Fig. 2f), consistent with the ATACseq and transcriptomic data.

Fig. 2g shows the distribution of M2_IL4_-specific, M2_Dex_-specific, or shared peaks when split into hyper-(red) or hypoacetylated (blue) subsets. Reflecting the trend seen in the ATACseq data, many more of shared sites were hypoacetylated. Importantly, putative shared enhancers included those near ‘hallmark’ M2 genes (*Klf9*, *Klf4*, and *Chil3*; underlined), whereas M1-like genes (*Irak3*, *Card11*, *Slamf9* and *CD300e*) lost associated H3K27ac marks (Fig. 2h). At the same time, other peaks associated with M2 genes gained H3K27ac marks in a signal-specific manner (*Pparg* and *Egr2* for M2_IL4_ and *Tsc22d3* and *Ms4a4a* for M2_Dex_), suggesting that the distinctions in chromatin landscape between the two populations correlate with population-specific transcription profiles. We validated hyperacetylation at the enhancer regions of key M2 target genes *Arg1*, *Klf9*, and *Chil3* by ChIP-qPCR (Extended Data Fig. 2c).

To focus specifically on enhancer elements potentially controlling transcriptional output in M2_IL4_ and M2_Dex_ populations, we excluded 1,650 differential acetylation sites which overlapped transcription start sites (TSS: –300/+200, Fig. 2i). The analysis of the remaining 12,211 peaks revealed an even greater overlap between M2_IL4_ and M2_Dex_: 42% of all M2_Dex_ differential enhancers overlapped with those seen in the IL4-polarized macrophages. We then stratified signal-specific and shared enhancers by the direction of regulation relative to those in M0 (Fig. 2j). Interestingly, and further corroborating unfiltered acetylation data in Fig. 2g, among the M2_IL4_:M2_Dex_ common enhancers, we detected over twice as many hypoacetylated enhancers as the hyperacetylated ones (827 vs. 390).

To better understand upward and downward changes in H3K27 acetylation, we performed rank shift analysis conditional on treatment. We ranked combined M2_Dex_ and M2_IL4_ differential peaks by mean H3K27ac signal (inverse log2CPM) in M0 and M2_IL4_ macrophages and plotted peak rank in M0 vs. the difference in peak ranks between M2_IL4_ and M0 (Extended Fig. 2d). We used LOESS regression to visualize the relationships between acetylation strength in M0 and signal change in M2_IL4_ (green line). Although the rank gain in M2_IL4_ macrophages is larger in magnitude, a larger fraction of peaks lost acetylation signal upon polarization.

Interestingly, peaks with large rank change were associated with genes with large expression change in M2_IL4_ (Extended Fig. 2d, right panel). Finally, as also indicated in Fig. 2f-g and i-j, and consistent with transcriptomic and chromatin accessibility data, most shared differential peaks in M2_IL4_ and M2_Dex_ change in the same direction, suggesting that genome-wide convergence of IL4– and GC-dependent programming occurs at multiple regulatory levels.

### Epigenomic and transcriptomic programming are tightly correlated in both M2_IL4_ and M2_GC_ macrophage populations

The expression of M2_IL4_– and M2_Dex_-specific (e.g., *Chil4* and *Hif3a*) and the M2_IL4_:M2_Dex_ shared (e.g., *Klf9*) genes tracked closely with changes in chromatin accessibility and H3K27 acetylation (Fig. 3a), which prompted us to systematically examine genome-wide relationships between gene expression and chromatin state during macrophage polarization by IL4 and GC. We therefore selected all differential ATACseq and H3K27ac ChIPseq peaks in –20 Kb to +20 Kb windows centered on TSS. We first stratified genes (TSS windows) by the gene status in our RNAseq (Fig. 1a) into differentially expressed in M2_IL4_ and/or M2_Dex_ vs. non-differential. We subsampled non-differential genes to create random sets of the same size as differential gene sets (n=10). For each gene in either group, we determined the number of associated ATACseq or H3K27ac peaks and selected peaks with the largest (positive or negative) logFC from M0 to either M2_IL4_ or M2_Dex_. We observed that polarization-driven changes in both ATACseq and H3K27ac peaks, associated with differential genes, corresponded to the direction of change in gene expression (Fig. 3b). The upregulated genes in M2_IL4_ and M2_Dex_ populations showed larger signal gain in ATACseq and H3K27ac peaks compared to non-differential genes, whereas downregulated genes exhibited greater loss of ATACseq and H3K27ac ChIPseq signal (Fig. 3b).

**Fig. 3.**
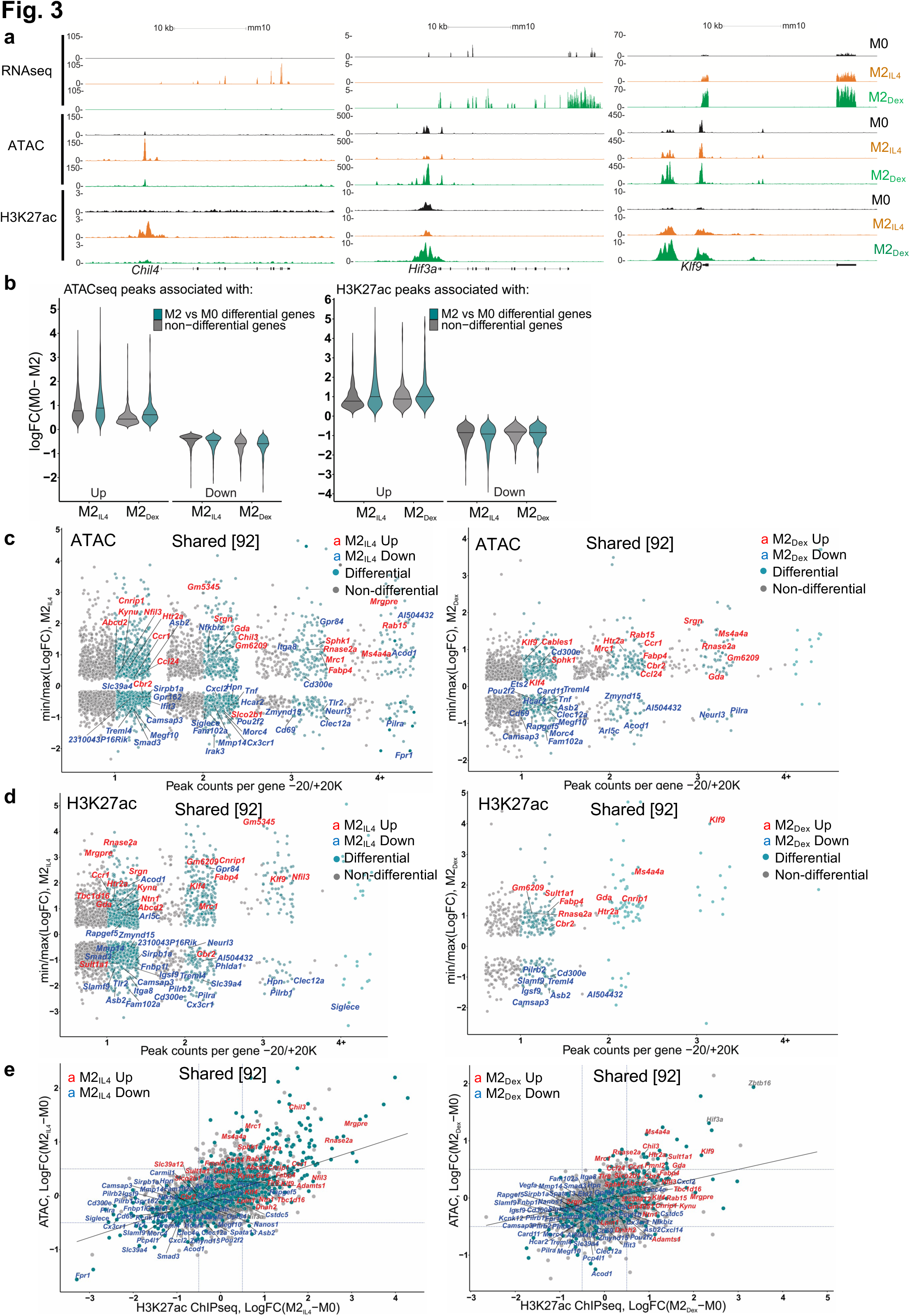
| M2Dex and M2_IL4_ have both distinct and shared *de novo* enhancers as demonstrated by H3K27ac ChIPseq. **a**, Correlation between RNAseq, ATACseq and H3K27ac ChIPseq signals in selected M2_IL4_-specific, M2_Dex_-specific and shared genes. **b,** Genome-wide changes in up-or downregulated differential ATACseq (left) and H3K27ac ChIPseq (right) peaks located within –20K +20K window centered on the TSS in M2_IL4_ and M2_Dex_, as indicated. **c-d,** Comparison of TSS window-associated ATACseq (**c**) or H3K27ac ChIPseq (**d**) peaks in differential (teal) vs. non-differential (gray) genes in M2_IL4_ (left panels) and M2_Dex_ (right panels), as indicated, stratified by the number of gene-associated peaks. Only peaks with the largest positive (max) or negative (min) changes relative to M0 are shown. A subset of 92 shared up-(red) and downregulated (blue) DEGs identified by RNAseq (Fig. 1) are labeled in each of the 4 panels. **e,** Correlation between normalized H3K27ac ChIPseq and ATACseq signals in the TSS-proximal –1K +1K windows for differential (teal) and non-differential (gray) genes. A subset of 92 shared upregulated (red) and downregulated (blue) DEGs identified by RNAseq (Fig. 1) are labeled.

We further stratified genes (TSS windows) by the number of associated ATACseq and H3K27ac ChIPseq peaks and plotted them against largest / smallest normalized peak signals for the corresponding genes in M2_IL4_ and M2_Dex_ populations. Notably, both M2_IL4_ (Fig. 3c and d, Extended Data Fig 3a and b, left) and M2_Dex_ (Fig. 3c and d, Extended Data Fig. 3a and b, right) DEGs (teal) were overwhelmingly associated with the larger number of dynamic ATACseq and H3K27ac peaks compared to non-differential genes (grey). Indeed, many of the M2_IL4_:M2_Dex_-shared (92) as well as M2_IL4_ (587)– and M2_Dex_ (174)-specific DEGs identified by RNAseq (Fig. 1a) including shared upregulated (Fig. 3c and d: *Ms4a4a*, *Mrc1, Ccr1*, *Klf9* and *Klf4*), shared downregulated (Fig. 3c and d: *Cd69*, *Tnf*, *Igsf9*), M2_IL4_-unique (Extended Data Fig. 3a and b: *Itgax, Msx3, Il1r1*) and M2_Dex_-unique (Extended Data Fig. 3a and b: *Fkbp5, Pi3ip1, Il15ra, Tsc22d3*) were associated with dynamic ATACseq and H3K27ac peaks. Although not all DEGs were associated with differential ATACseq and H3K27ac peaks, these data reveal a remarkable level of convergence between polarization-driven chromatin accessibility and H3K27 acetylation vs. transcriptomic programming.

To compare M2 polarization-driven chromatin and transcriptional changes directly, we selected all 9,323 genes expressed in any condition in our RNAseq experiment and counted the reads overlapping TSS-centered –1Kb /+1Kb regions in both ATACseq and H3K27ac ChIPseq experiments. Using thus generated TSS-proximal count matrices, we performed separate differential, relative to M0, analyses in M2_IL4_ and M2_Dex_ for ATACseq and H3K27ac to identify genes whose TSS-proximal regions undergo changes in accessibility and acetylation status during M2 polarization. This analysis revealed a striking correlation between chromatin accessibility, H3K27 acetylation and gene expression in both M2_IL4_ and M2_Dex_ (Fig. 3e and Extended Data Fig. 3c-f). Indeed, M2-like M2_IL4_:M2_Dex_-shared (*Klf9*, *Mrc1*, and *Nfil3*) and signal-specific (*Arg1* and *Chil4* for M2_IL4_ and *Zbtb16* and *Hif3a* for M2_Dex_) genes gained H3K27ac mark that also corresponded to a more open chromatin, whereas M1-like genes (*Tnf*, *Ifit3*, and *Pilra*) lost both ATACseq peaks and associated H3K27ac mark (Fig. 3E and Extended Data Fig. 3c-f). Furthermore, the Ingenuity Pathway Analysis (QIAGEN) of all M2_IL4_ and M2_Dex_ DEGs associated with changes in both the ATACseq and H3K27ac signals, revealed enrichment of pathways linked to known M2 functions such as cellular movement, immune cell trafficking, and cellular function and maintenance (Extended Data Fig. 3g; Extended Data Table 2). Collectively, these analyses suggest that transcriptomic changes that occur during macrophage polarization by IL4 and Dex are tightly correlated with changes in enhancer landscape and overall chromatin accessibility.

### Inducible co-localization of GR, KLF4 and transcriptional coregulator GRIP1 across the genome

IL4 and GCs act through distinct receptors in unrelated protein families and initiate signaling events that have little in common. Therefore, we reasoned that the striking overlap in the transcriptomic profiles of M2_IL4_ and M2_GC_ is generated through common utilization of cofactors at the level of downstream effector transcription complexes. GR is a classic ligand-dependent nuclear receptor which relies on the p160 family of coregulators – NCoA1, 2 and 3 ^34^ – of which NCoA2 (GRIP1/TIF2/SRC2 – hereafter GRIP1) has been implicated both in GR-driven gene activation and repression in various cell types including macrophages ^20, 35, 36^. Interestingly, we previously described an unexpected interaction between GRIP1 and KLF4, and their co-recruitment to *Arg1* and *Klf4* genes in IL4-treated macrophages ^37^. We therefore asked whether GRIP1 functioned globally to integrate GC– and IL4-driven M2 polarization. To begin to dissect genome-wide relationships between GR, KLF4 and GRIP1, we first performed GR ChIPseq in M0, M2_IL4_, M2_Dex_ and M2_Cort_ populations. As expected, GR peaks in M2_Dex_ and M2_Cort_ fully overlapped, and GR binding was strictly GC ligand-dependent with no specific peaks detected in M2_IL4_ (Fig. 4a and Extended Data Fig. 4a), including near genes that were transcriptionally activated in response to both IL4 and GCs (e.g., *Klf9* in Fig. 4c). Transcription factor binding motif enrichment analysis showed that the dominant motif in both M2_Dex_ and M2_Cort_ GR cistromes was the GRE/ARE (NR3C nuclear receptor binding motif; Extended Data Fig. 4b).

**Fig. 4.**
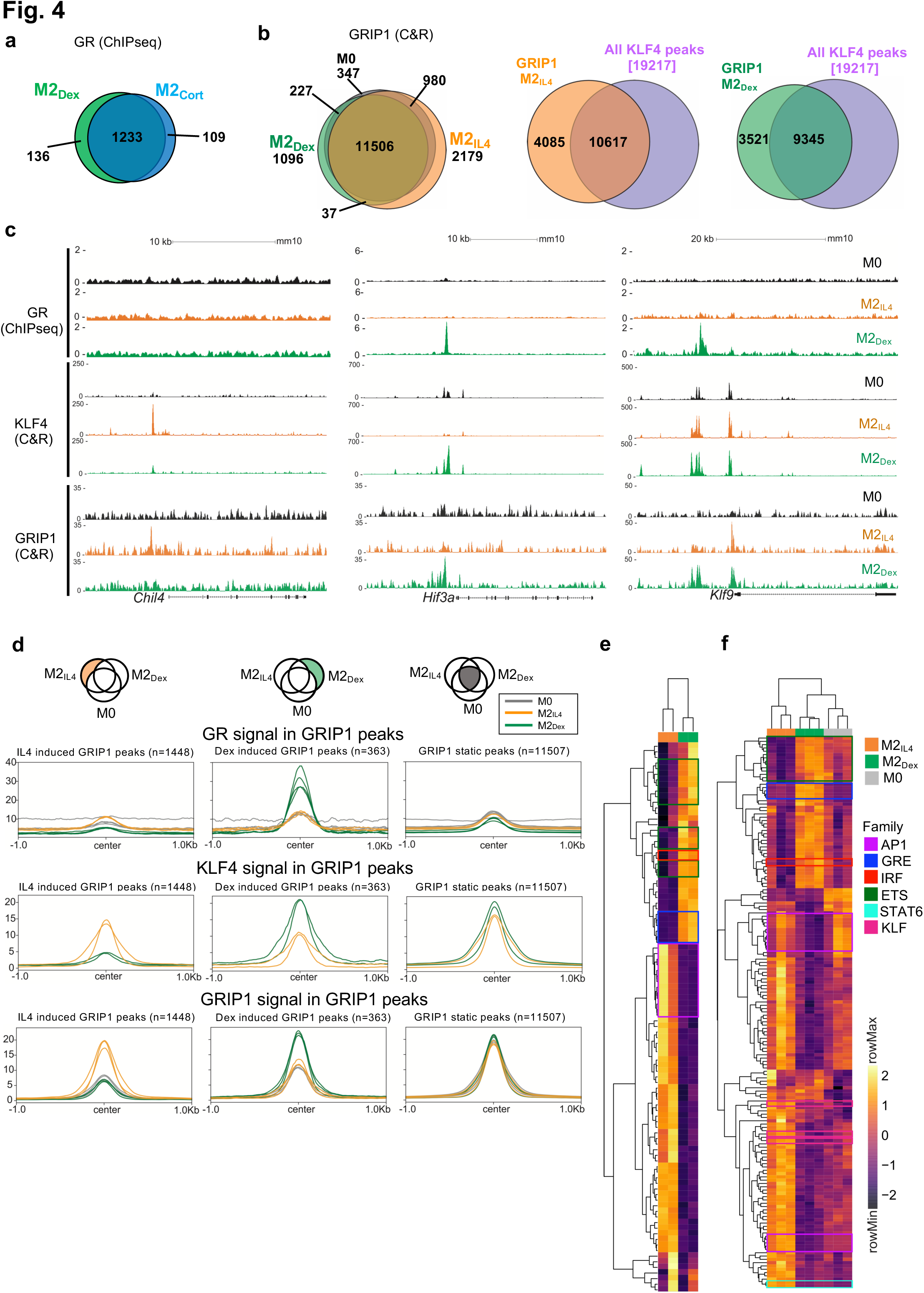
| GR, KLF4 and GRIP1 genome-wide binding in differentially polarized macrophage populations. **a**, Venn diagram shows the numbers and overlap of GR ChIPseq peaks in M2_Dex_ and M2_Cort_ macrophages relative to M0 baseline (n=4; FC ≥ 2, FDR <0.05). **b,** Venn diagram shows the numbers and overlap of (left) GRIP1 CUT&RUN peaks in M0, M2_IL4_ and M2_Dex_ populations (n=3; FC ≥ 1.5, FDR <0.05) or (middle and right) GRIP1 in M2_IL4_ and all KLF4 peaks (n=2) or GRIP1 in M2_Dex_ and all KLF4 peaks, as indicated. **c,** GR (ChIPseq), KLF4 (CUT&RUN) and GRIP1 (CUT&RUN) read density distribution over *Chil4*, *Hif3a* and *Klf9* loci in M0, M2_IL4_ and M2_Dex_. **d,** Average profiles of GR, KLF4 and GRIP1 signal from each macrophage population centered on GRIP1 CUT&RUN M2_IL4_-(left), M2_Dex_-(middle) specific or invariant ‘static’ (right) peaks (n=3; FC ≥ 2, FDR <0.05). **e-f,** Transcription factor binding motif enrichment Z-scores in KLF4 (**e**) and GRIP1 (**f**) peak subsets in each macrophage population (M2_IL4_=orange, M2_Dex_=green, M0=grey, as indicated) were determined in chromVAR from the HOMER list of motifs. Motifs with high variability among the samples (p_adj_ <= 1*10^-6^) were plotted and indicated motif families are highlighted in boxes.

We then employed CUT&RUN to assess genomic occupancy of GRIP1 and KLF4 in differentially polarized M2 macrophage populations. GRIP1 bound extensively across the genome in M0 macrophages, with the majority of peaks maintained during polarization (Fig. 4b, left; 347 present in M0 only of the 16,372 total GRIP1 peaks). Polarization induced thousands of additional GRIP1 binding sites (Fig. 4b, left; 2,216 for M2_IL4_ and 1,133 for M2_Dex_ with 37 peaks overlapping). We generated a KLF4 peak set from peaks present in either polarized population (19,217 peaks) to assess its overlap with the GRIP1 binding. Strikingly, the vast majority of GRIP1 peaks, 10,617 of total 14,702 in M2_IL4_ and 9,345 of 12,866 in M2_Dex_, overlapped with at least one KLF4 peak (Fig. 4b, right). Although the total putative binding sites for GRIP1 and KLF4 generated by CUT&RUN considerably exceeded the ChIPseq dataset for GR, we still found that 44% of KLF4 intersections and 27% of GR intersections involve GRIP1 peaks (Extended Data Fig. 4c).

Given the dramatic overlap between GRIP1 occupancy and that of the two transcription factors driving the M2 phenotype in our differentially polarized macrophages, we evaluated binding of all three proteins near M2_IL4_-specific *Chil4*, M2_Dex_-specific *Hif3a* and shared *Klf9* in all three populations. No GR or GRIP1 peaks and minimal KLF4 binding were detected near these genes in M0 macrophages, demonstrating the signal specificity of these binding events (Fig. 4c). As expected, in M2_Dex_, GRIP1 was recruited to GR-bound sites of *Hif3a* and *Klf9* – the same sites that gained H3K27ac mark and peaks of chromatin accessibility (Fig. 3a). Also consistent with chromatin activation, in M2_IL4_, KLF4 was recruited to the acetylated *Chil4* enhancer, together with GRIP1 (Fig. 4c, orange tracks for KLF4 and GRIP1). Similarly, KLF4 also bound to two sites upstream of *Klf9* and the promoter-proximal site was co-occupied by GRIP1. Of note, the upstream KLF4 peak in M2_IL4_ appeared to co-localize with the GR peak in M2_Dex_, perhaps pointing to the adjacent binding sites for the two factors. Surprisingly, KLF4 was also recruited in Dex-polarized macrophages (Fig. 4c, green tracks for KLF4) to the GR:GRIP1-co-occupied sites of *Hif3a* and *Klf9*, as well as promoter-proximal *Klf9* site not bound by GR, yet still recruiting GRIP1. These observations suggest that, unlike strictly GC-dependent actions of GR, KLF4 and GRIP1 occupancy is much more dynamic and ‘permissive’, and that upregulation of KLF4 expression by both signals, combined with GRIP1 binding by both GR and KLF4 create the chromatin environment that facilitates the shared epigenetic and transcriptional programs induced by the two pathways.

Consistent with this idea, when quantified in condition-specific GRIP1 peak subsets, both KLF4 and GRIP1 signals were stimulation-specific for both M2_IL4_ and M2_Dex_, and GR signal was only enriched in M2_Dex_ macrophages and was present in Dex-induced GRIP1 peaks only (Fig. 4d, left and middle columns). In contrast, no obvious bias for KLF4 or GR enrichment in any population was seen in ‘static’ GRIP1 peaks (Fig. 4d, right column).

We then focused on transcription factor motifs enriched in the KLF4 and GRIP1 cistromes. As expected, KLF family binding motifs were broadly over-represented in the KLF4 peaks independent of stimulus origin (Extended Data Fig. 4d). Importantly, binding sites for distinct families of transcription factors were enriched in different macrophage populations. In both KLF4 and GRIP1 cistromes, the NR3C/GRE motif was enriched specifically in GC-polarized macrophages (Fig. 4e-f, Extended Data Fig. 4e). This finding is particularly interesting for the KLF4 cistrome, as it further underscores the co-localization of KLF4 and GR binding sites. A similar pattern was seen for lineage-determining ETS family and, perhaps surprisingly, IRF (Fig 4e-f). Conversely, the KLF motif was over-represented in M2_IL4_ and to a lesser degree in M2_Dex_ GRIP1 peaks and colocalized with GREs in GRIP1 M2_Dex_ peaks (Fig 4f, Extended Data Fig. 4e). Interestingly, a binding motif for STAT6 – a direct downstream effector of IL4 signaling – was enriched selectively in M2_IL4_ GRIP1 peak subsets and was found in M2_Dex_ GRIP1 peaks containing both GR and KLF4 motifs (Fig. 4f, Extended Data Fig. 4e). Together, these experiments reveal broad, yet specific co-localization of GR:GRIP1 and KLF4:GRIP1 transcription complexes across the genome in differentially polarized macrophage populations.

### GRIP1 facilitates the establishment of M2_IL4_ and M2_GC_ transcriptomes

Given that GRIP1 is a transcriptional coregulator that colocalized with KLF4 and GR at regulatory regions associated with many M2-enriched genes, we examined its contribution to IL4– and GC-induced transcriptional programming. We assessed gene expression in M0, M2_IL4_, M2_Dex_ and M2_Cort_ derived from mice of two genotypes: WT (LysM-Cre^+/+^; GRIP1^wt/wt^) and GRIP1 cKO (LysM-Cre^+/+^; GRIP1^fl/fl^) lacking GRIP1 in the myeloid lineage ^38^ and, hence, in all four polarized macrophage populations (Fig. 5a). RT-qPCR revealed (Fig. 5b) that in GRIP1-deficient macrophages, the induction of individual candidate M2 genes was dramatically reduced irrespective of whether they were M2_IL4_-specific (top), M2_GC_-specific (middle) or shared (bottom).

**Fig. 5.**
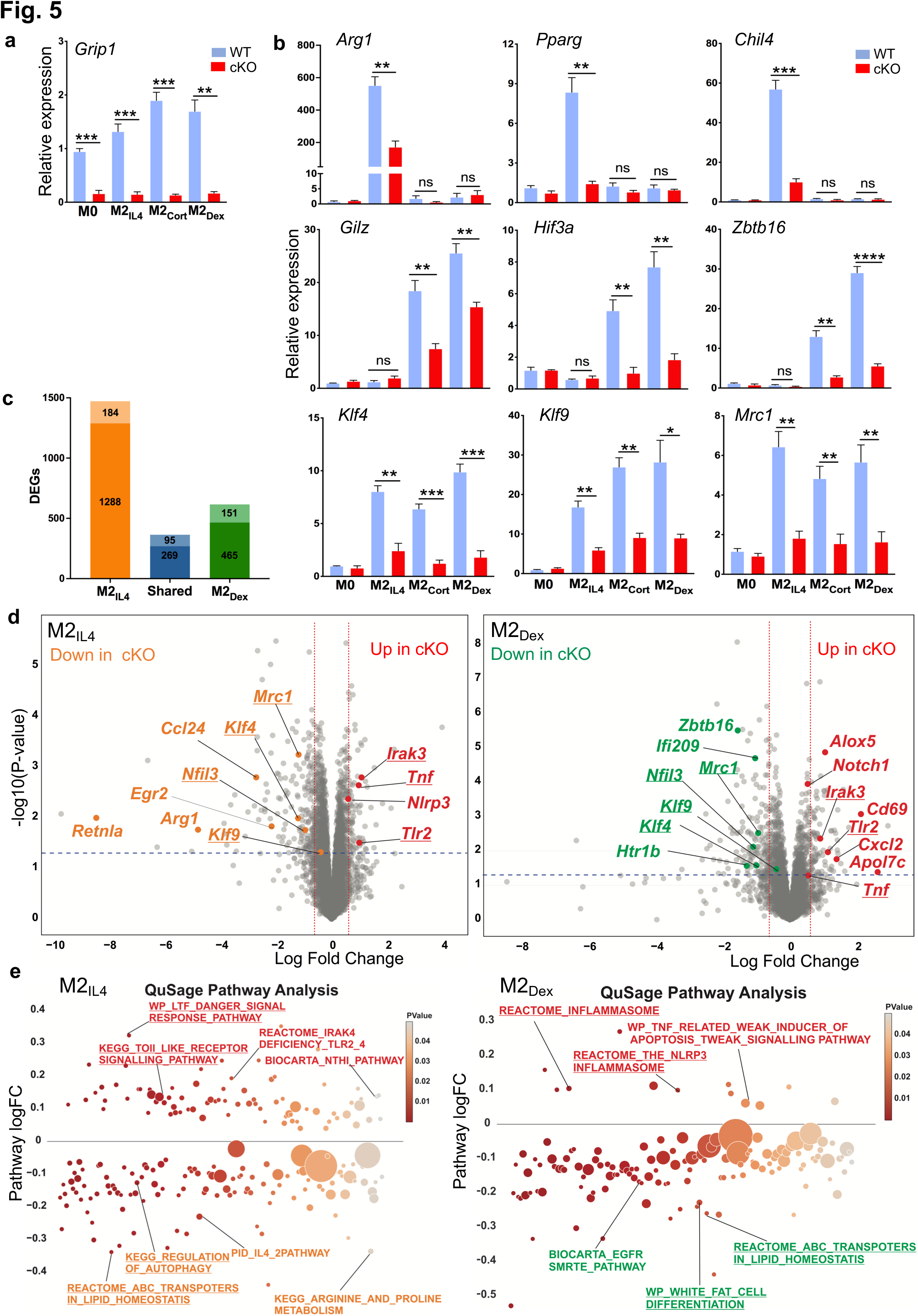
| GRIP1 is required for the establishment of M2_IL4_ and M2_GC_ transcriptomes. **a-b**, Expression of (**a**) GRIP1 or (**b**) IL4-(top) GC-(middle) or shared (bottom) M2 target genes in WT and GRIP1 cKO M0, M2_IL4_, M2_Cort_, and M2_Dex_ populations as assessed by RT-qPCR. Average relative expression of each transcript in WT M0 is arbitrarily set to 1. n=4; Student’s t-test; *p < 0.05, **p < 0.01, ***p < 0.001, ****p < 0.0001, ns, non-significant. Error bars are SEM. **c,** Stacked bar plot shows the number of DEGs in GRIP1 cKO vs. WT as a fraction of WT M2_IL4_, M2_Dex_, or shared DEGs relative to M0 (RNAseq, WT: n=3, cKO: n=4). **d,** Volcano plot shows DEGs in GRIP1 cKO vs. WT in M2_IL4_ (left) and M2_Dex_ (right) relative to M0 of each genotype. (FC≥1.5, unadjusted p-value<0.05). Selected inflammation-related genes expressed at higher levels in the cKO M2_IL4_ and M2_Dex_ relative to WT are highlighted in red. Key M2 target genes downregulated in the cKO are highlighted in both M2_IL4_ (orange) and M2_Dex_ (green). Examples of the shared M2_IL4_:M2_Dex_ GRIP1 target genes are underlined. **e,** Differentially regulated pathways (unadjusted p < 0.01) identified using QuSAGE with MsigDB canonical pathways subset (Broad Institute) are shown for GRIP1 cKO vs. WT in M2_IL4_ (left) and M2_Dex_ (right). Select pathways are labeled (upregulated, red; downregulated, orange for IL4 and green for Dex); M2_IL4_:M2_Dex_-shared pathways are underlined. Circle size is proportional to the number of genes in the pathway and color signifies p-value.

To obtain a more accurate assessment of the genome-wide impact of GRIP1 deficiency and focus on genes that lose most of their transcriptional response to polarizing signals, we adjusted the RNAseq cut-off FC from 2-fold (log2FC≥1) to 1.5-fold (log2FC≥0.6). Under these conditions, we identified 1,288 and 465 DEGs, relative to M0, in WT M2_IL4_ and WT M2_Dex_, respectively, and 269 genes that were shared (Fig. 5c). Loss of GRIP1 attenuated the regulation of 184 (or 13%) in the M2_IL4_ transcriptome. A much greater fraction – 151 genes (or 25%) – in the M2_Dex_ transcriptome was GRIP1-dependent. This is consistent with a well-established function of GRIP1 as a GR cofactor, implicated in both GR-mediated repression and activation *in vivo* ^20, 36^. Interestingly, the fraction of GRIP1-dependent genes was highest for the M2_IL4_:M2_Dex_-shared transcriptome: 95 (26%) of the shared DEGs required GRIP1 for their regulation.

The differential gene expression analysis revealed that GRIP1 contributes to the expression of key orchestrators of M2 programming in each population. A subset of GRIP1-regulated genes specific to M2_IL4_ included major transcription factors known to confer the M2_IL4_ state (e.g., *Egr2*), as well as other signature M2_IL4_ genes (*Arg1*, *Retnla*) (Fig. 5d, left), suggesting the functional significance of this gene-set despite its relatively modest size. Similarly, GRIP1 was required for the full induction of M2_GC_-specific genes, *e.g.*, *Htr1b* and *Zbtb16* (Fig. 5d right and Extended Data Fig. 5a). Importantly, many GRIP1-dependent genes including M2 signature genes *Klf4* and *Mrc1* were overlapping in IL4– and GC-polarized macrophages, consistent with a large portion of the shared M2_IL4_:M2_GC_ transcriptome being GRIP1-dependent (Fig. 5d and Extended Data Fig. 5a, underlined). Interestingly, the expression of a subset of genes related to the inflammatory M1 phenotype such as *Irak3*, *Tlr2*, and *Tnf* was higher in GRIP1 cKO relative to WT in M2_IL4_, M2_Dex_ and M2_Cort_ transcriptomes (Fig. 5d and Extended Data Fig. 5a), indicating that in all three cases, GRIP1 contributed to the repression of these genes.

GRIP1-regulated genes form distinct pathways along the spectrum of macrophage polarization, encompassing both the homeostatic and pro-inflammatory functions. QuSAGE analysis demonstrated dampening of the M2 pathways in M2_IL4_ and M2_GC_ in the GRIP1 cKO (Fig. 5e and Extended Data Fig. 5b; Extended Data Table 3). For the M2_IL4_ population, the Arginine and Proline Metabolism (hallmark of metabolic activity of M2_IL4_) and Regulation of Autophagy (associated with M2 phagocytic activity) were downregulated (Fig. 5e, left). The GC-specific EGFR SMRTE_Pathway was downregulated in the cKO M2_GC_ macrophages (Fig. 5e, right and Extended Data Fig. 5b). In contrast, the pathways associated with inflammatory signatures were upregulated in all three cKO populations (Fig. 5e and Extended Data Fig. 5b).

Because both IL4 and Dex induced dramatic changes in macrophage chromatin accessibility during M2 polarization (Fig. 2a), we asked whether GRIP1 contributes to alteration in chromatin environment in M2_IL4_ and/or M2_Dex_. We performed ATACseq in WT and GRIP1 cKO which showed that overall peak distribution across the genome was similar in WT and cKO in both M2_IL4_ and M2_Dex_ populations (Extended Data Fig. 5c). Genome-wide comparison of ATACseq peaks in M2_IL4_ and M2_Dex_ between the two genotypes yielded relatively modest subsets of uniquely IL4– and Dex-regulated peaks (329 and 457, respectively) and 400 shared differential peaks that were affected by loss of GRIP1 (Extended Data Fig. 5d). Among those, 395 and 414 sites were ‘upregulated’ (failed to close) in cKO M2_IL4_ and cKO M2_Dex_, respectively, compared to matching WT populations; conversely, 334 and 443 peaks, respectively, were ‘downregulated’ (failed to open) in cKO M2_IL4_ and cKO M2_Dex_ (Extended Data Fig. 5e). Collectively, these data suggest that GRIP1 is required for establishing the transcriptional state in M2_IL4_ and M2_Dex_ populations; its exact contribution to *de novo* IL4– and GC-driven changes in chromatin accessibility remain to be determined.

### GRIP1 contributes to the phagocytic activity of M2_IL4_ and M2_GC_ *in vitro* and tissue healing in the mouse dextran sulfate sodium (DSS)-induced colitis *in vivo*

Phagocytic activity is a hallmark function of M2 macrophages, needed for engulfing apoptotic cells and cell debris ^39^. We set out to test if GRIP1 is necessary for phagocytosis in M2_IL4,_ M2_Dex_ and M2_Cort_. When exposed to bioparticles that fluoresce when internalized into the acidic phagosome, WT macrophages polarized with either IL4 or GCs exhibited approximately 20% relative increase of phagocytosis *in vitro* (Fig. 6a). Interestingly, GRIP1 deletion dramatically reduced their phagocytic activity compared to the WT. Indeed, M2_IL4_ GRIP1 cKO macrophages reverted to the baseline phagocytic activity (equal to that of M0), whereas M2_GC_ show an attenuated induction. We conclude that GRIP1 contributes to the upregulation of phagocytic activity in both M2_IL4_ and M2_GC_.

**Fig. 6.**
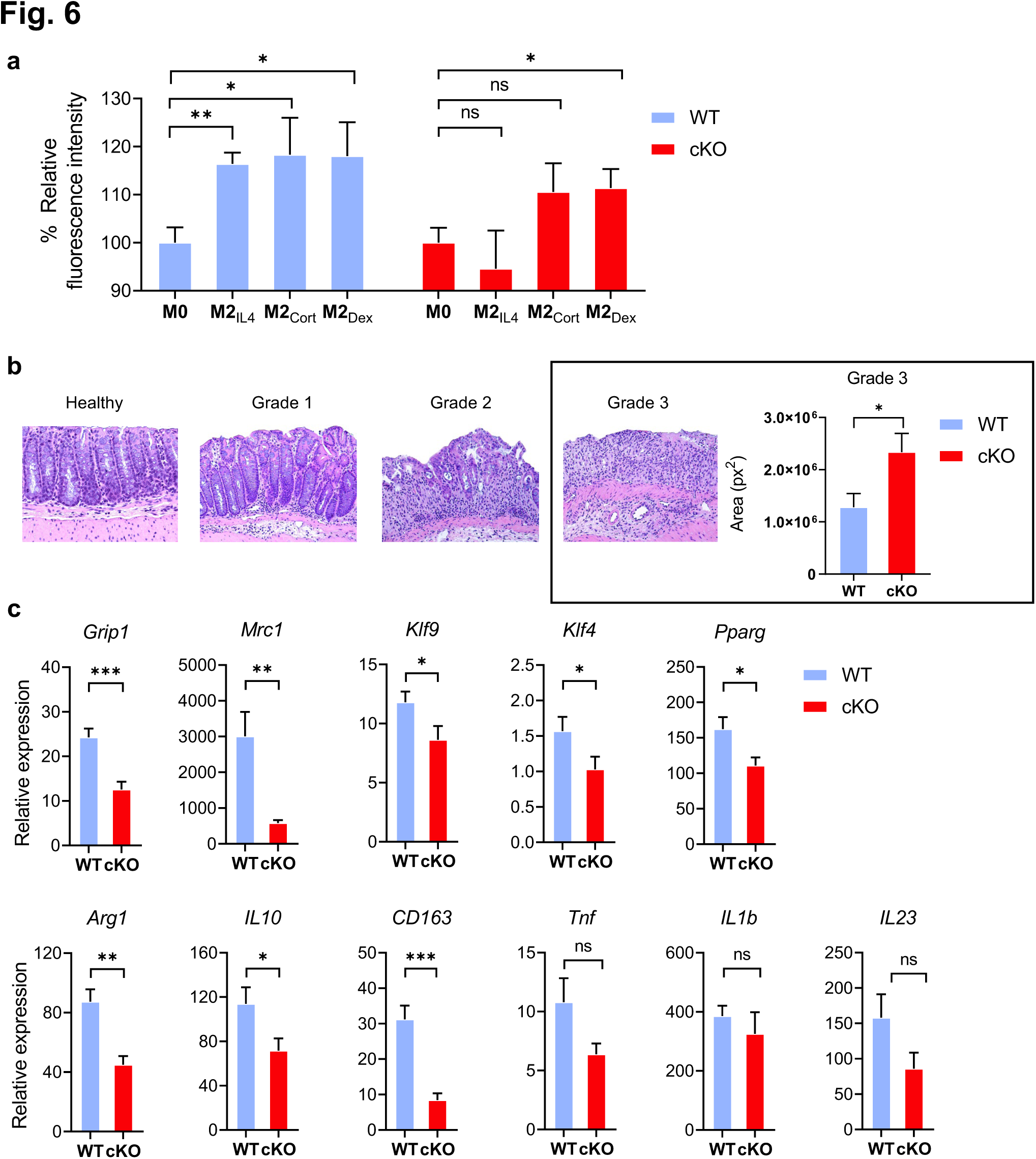
| Macrophage GRIP1 contributes to phagocytic activity of M2_IL4_ and M2_GC_ *in vitro* and tissue healing *in vivo*. **a**, Relative fluorescence intensity of WT (set to 100%) and GRIP1-cKO M2_IL4_, M2_Cort_ and M2_Dex_ relative to that in M0 after exposure to pHrodo Green *S. aureus* BioParticles (see Methods). Shown are mean values and error bars are SEM (n=3, student t-test). **b,** Histopathological changes assessed by H&E staining of colons from WT and cKO mice 2 wks after initiating DSS-induced colitis. Grade 1, 2 and 3 are defined in Methods. Plotted is the total area representing grade 3 for colon specimens of WT and GRIP1-cKO (n=4, student t-test). **c,** The expression of indicated genes was measured by RT-qPCR of total RNA from colons of WT and cKO mice. The expression of each gene was normalized to β-actin, and then fold difference to baseline, non-treated (calibrant) value calculated for WT or cKO. Shown are mean values and error bars are SEM (WT: n=10-15, cKO: n=6-8; student t-test).

We next tested whether GRIP1 promotes the homeostatic tissue-repair macrophage phenotype *in vivo* in an acute inflammatory setting. We chose the DSS-induced colitis model, since virtually all macrophages in the colon are derived from monocytic progenitors ^40^, rendering the model consistent with our reliance on BMDM *in vitro*. In addition, over the duration of the model, both inflammatory and tissue-repair macrophages function simultaneously, circumventing complexities of interpreting changes in the sequential recruitment of macrophage populations. When exposed to 2% DSS-containing water for 10-14 days, both WT and GRIP1 cKO mice displayed characteristic histopathological features of colitis. The histopathological changes were graded as: (1) well defined crypts and villi with preserved mucus secretion and a mild lymphocytic inflammation; (2) the villi are short and distorted, crypts and goblet cells are diminished, apparent hyperplasia of glandular and superficial epithelium; (3) loss of crypts, ulceration of mucosa, severe inflammation and heavy infiltration of lymphocytes, macrophages and granulocytes (Fig. 6b). Importantly, GRIP1 cKO mice developed significantly greater tissue destruction as total area occupied by the most severe grade 3 damage was doubled over that seen in the WT (Fig. 6b). Consistently, gene expression analysis of total colonic RNA revealed that expression of many typical M2 genes including *Klf4, Klf9, Mrc1* and *Arg1* was significantly attenuated in the GRIP1 cKO mice relative to their WT counterparts (Fig. 6c), resembling the results of GRIP1 deficiency *in vitro* in the M2_IL4_ and M2_GC_ populations (Fig. 5b and c).

Unexpectedly, the inflammatory genes known to be involved in colitis progression (*Tnf, Il1b, Il23*) were expressed at similar levels in both genotypes (Fig. 6c). Body weight, colon length and weight were similar between genotypes (Extended Data Fig. 6). We conclude that the greater susceptibility of GRIP1 cKO mice to DSS-mediated colonic tissue destruction, at least at the end-point of the disease, is due to the failure of cKO macrophages to repair the damage, rather than increased inflammation in the colon. Thus, GRIP1 contribution to extensive transcriptomic rewiring of macrophages during the establishment of their homeostatic state is required for their adequate tissue repair properties *in vivo*.

## Discussion

Phenotypic similarity between macrophages exposed to type-2 cytokines and those treated with GCs, including their phagocytic activities ^41, 42^, wound healing and tissue repair functions ^43, 44^, has been recognized for a long time in both human and murine systems. This was further supported by early microarray studies which juxtaposed the transcriptomic features of these *in vitro* produced ‘M2’ populations to more traditionally inflammatory ‘M1’ macrophages ^19, 45^. The binary model of macrophage activation was later challenged in a seminal study by Xue et al, which highlighted the remarkable plasticity of macrophage responses, whereby dozens of stimuli yielded a wide spectrum of transcriptomic states ^9, 46^. Nonetheless, macrophages elicited by type-2 cytokines and GCs still populated the same ‘pole’ of the spectrum, farthest away from that elicited by inflammatory stimuli such as IFNγ and LPS ^19^. Here, we aimed to understand whether and how two highly divergent biological signals – IL4 and GC – trigger converging polarization programs, the epigenomic wiring that enforces these programs and the mechanisms that underlie their establishment.

We focused on stable changes that occur by 24 h of IL4/GC exposure. We reasoned that unlike rapid transcriptional responses of genes in open chromatin, often preloaded with fully engaged, initiated RNA Polymerase II ^21, 47, 48^, epigenomic changes accumulate more slowly and are more indicative of the macrophage transition into a novel phenotypic state. Indeed, a recent study classified enhancers in IL4-treated macrophages into ‘early transient’, ‘early sustained’ and ‘late’ ^14^. All three groups exhibited p300 and H3K27ac gain, detectable at 24 h of IL4 exposure only for the sustained and late enhancers. Interestingly, after filtering out promoters from the H3K27ac dataset, we obtained 5248 IL4-induced enhancers, comparable to the total of 7309 sustained and late enhancers reported by Bence et al. Furthermore, that study showed that a subset of the activated enhancers are occupied by the histone reader BRD4 and elongating RNA Polymerase II. Thus, histone bookmarking and active enhancer transcription both signify the establishment of the M2_IL4_ state. In addition, the transcription factor EGR2 was proposed to be required for the acquisition of the late enhancer networks ^10, 14^ – and we detected EGR2 upregulation, the enrichment of EGR2 binding sites in the opening chromatin and activation of *Egr2*-associated enhancers specifically in M2_IL4_. Interestingly, STAT6 motif was overrepresented among the open motifs in both M2_IL4_ and M2_Dex_. This was unexpected in light of earlier observations that recruitment of STAT6 – a direct effector of IL4 signaling – largely fades away by 24 h, while EGR2 remains bound ^14^. This may indicate that the chromatin changes near STAT6 binding sites persist long beyond transcription factor occupancy.

Unlike these exciting developments in our understanding of the epigenomic pathways driving macrophage responses to IL4, little is known about the long-term impact of GCs on macrophage programming. GC signaling elicits profound anti-inflammatory effects *in vivo*, which made synthetic GCs some of the top prescribed medications worldwide ^49^. In the context of macrophages, therapeutic efficacy of these molecules has been historically attributed to broad transcriptional repression of genes encoding inflammatory cytokines, chemokines, tissue degrading enzymes and other mediators of inflammation and tissue destruction ^50^. Indeed, many pro-inflammatory genes were downregulated in our M2_Dex_ and hundreds of associated chromatin sites were losing accessibility and the H3K27ac mark. Surprisingly, however, an overall greater number of genes and associated enhancers, including those with established M2-related functions (e.g., such as *Cebpb* ^51^, *Jdp2* ^52^, *NOTCH4* ^53^ and *MerTK* ^54^) were uniquely activated in M2_Dex_. In motif enrichment analysis, the GRE and the binding site for CEBP – an established GR-cooperating factor ^55^ were not only over-represented in open peaks in the M2_Dex_, but were more enriched in the Dex-polarized population relative to M0, indicating that many of the associated genes were direct transcriptional targets of GR – an observation further supported by GR ChIPseq.

Perhaps a larger surprise was the level of convergence between the M2_IL4_ and M2_GC_ populations, both in their chromatin states and their transcriptomes. Out of all the DEGs, acetylated enhancers and sites of opening chromatin in M2_Dex_, 30-50% overlapped with those regulated by IL4 and included hallmark M2 genes that confer the homeostatic macrophage phenotype, e.g., *Mrc1, Chil3* and *Nfil3*. In addition, we report a direct correlation between the transcriptomic and epigenomic changes in a 3-way comparison, which held true for both shared and signal-specific DEGs in each population. This suggests that the core M2 transcriptome is relatively stable with the transcriptomic changes likely to persist for extended periods of time.

Some population-specific DEGs orchestrate population-specific functions. *Arg1*, encoding arginase 1, drives the hydrolysis of arginine to urea and ornithine, thus promoting polyamine synthesis and decreasing nitric oxide production (a hallmark of M1-like phenotype) ^56^. Since *Arg1* upregulation was strictly IL4-specific, we anticipate that this metabolic shift does not occur in M2_Dex_. Conversely, several M2_GC_-specific DEGs are canonical GR targets, known to suppress inflammation (e.g, *Tsc22d3*) or regulate GR activity (e.g., *Fkbp5*) ^21^. At the same time, shared targets, like *Mrc1*, are directly involved in phagocytosis, consistent with the increase in phagocytic activity in both M2_IL4_ and M2_GC_ macrophages.

Given that IL4 and GCs signaling pathways are dramatically different, we reasoned that their functional convergence likely occurs at the level of chromatin and target gene regulation. Indeed, GR and KLF4 binding sites have been reported to colocalize throughout the genome ^30, 57^; KLF4 itself is upregulated by both IL4 and GC. Here we show that GRIP1, a nuclear receptor cofactor, mediates the transcriptional integration of the two states with 26% of the DEGs shared between M2_IL4_ and M2_Dex_ requiring GRIP1 for their regulation. We also observed a striking overlap between the GRIP1 and KLF4 cistromes which, unexpectedly, extended to GRIP1 peaks and DEGs specific to the M2_Dex_ population. Several mechanisms can potentially account for GRIP1-assisted KLF4 binding in GC-polarized macrophages. Given the proximity of GR and KLF binding sites, GRIP1 recruitment by GR may result in cofactor ‘spill-over’, enabling GRIP1 to bind and stabilize KLF4 on the DNA. Moreover, like other p160 family members, GRIP1 serves as a platform for many histone modifiers, such as acetyltransferases (HATs) CBP/p300 or arginine methyl transferases (CARM1) ^58, 59^, which would extend the regions of permissive chromatin to KLF4 binding sites. In fact, such cooperative recruitment may in turn further facilitate nucleosome remodeling ^60^. In addition, we envision such mechanisms to be broadly applicable to distant higher order 3D chromatin interactions, thus expanding the network of GRIP1-regulated genes.

One unexpected observation across different readouts was that the overlap between the M2_IL4_ and M2_Dex_ included more downregulated DEGs and inactivated enhancers than the upregulated ones. Thus far, all our comparisons were relative to unstimulated M0, whose characteristics may not fall into the center of the imaginary spectrum separating fully inflammatory macrophages from the homeostatic ‘M2’. Many actively expressed M2-like genes and enhancers are likely to already be acetylated. This baseline bias is also supported by the lack of prominent inflammatory signature among downregulated genes, relatively mild enrichment of the NFkB motif in deacetylated peaks and the absence of NFkB binding sites in the GR cistrome, which in M1-like macrophages are populated by GR through physical interactions with p65 ^21^.

Thus, our analysis underestimates the scale of transcriptional and epigenomic shift between the M1 and M2 populations, and likely, the true overlap between the M2_IL4_ and M2_Dex_ populations when they undergo this transition.

Finally, we acknowledge that the goal of this study – as well as its limitation – was to obtain a granular view of how a single stimulus, IL4 or GCs, rewires the epigenomic and transcriptional landscape. *In vivo*, macrophages encounter multiple signals, including type-1 vs. type-2 cytokines, TLR ligands, hormones or dead cells simultaneously, so their responses would reflect the compounded effects of disparate stimuli. Some of these scenarios have already been modeled *in vitro* ^39, 61, 62, 63^. In our transcriptomic studies, a small subset of genes (n=41) were regulated by IL4 and Dex in opposite directions, implying a potential antagonism, whereas the rest of the shared genes may display additive or synergistic relationship, as has been recently proposed ^64^. Our DSS-colitis model suggests that the defective tissue repair function of GRIP1 cKO macrophages rather than the uncontrolled inflammation was a dominant contributor to the phenotype at the end-stage of the disease. We do not, however, rule out the possibility that failure to contain inflammatory responses – which would be consistent both with our *in vitro* data and with established function of GRIP1 in the context of GC signaling ^20^ – aggravates earlier stages of disease. Future studies are required to untangle the complex and potentially timed relationships between macrophage responses and to determine the stability of their epigenomically programmed functional states. In particular, whether a pharmacological manipulation of GR or other key regulators can impart to a macrophage a long-lived non-inflammatory phenotype remains an important question for clinical care.

## Methods

### Mice

C57BL/6 mice (National Cancer Institute, Charles River Laboratories) and their transgenic derivatives were maintained in the Weill Cornell Animal Facility in compliance with the protocol approved by the Institutional Animal Care and Use Committee. Homozygous LysM-WT (LysM-Cre;GRIP^wt/wt^) and GRIP1-cKO (LysM-Cre;GRIP^fl/fl^) mice were generated previously^3737^. For all tissue and cell collection, mice were euthanized via CO2 inhalation.

C57BL/6 mice were used for ChIPseq experiments. WT and GRIP1-cKO mice were used for RT-qPCR, RNAseq, ATACseq, phagocytosis and colitis experiments. All mice used were 8-to10-week-old males.

### Cell culture

BMDM were prepared as previously described ^47^. Briefly, bone marrow was flushed from the femurs and tibiae of mice and differentiated for 5 days in the DMEM (1 g/l glucose) supplemented with 20% bovine serum and 10% L-cell conditioned media. Adherent cells were scraped, seeded into 150 mm or 6-well plates and polarized for 24 h with 10 ng/ml IL4, 500 nM Cort or 100 nM Dex unless otherwise indicated.

### RNA isolation and RT-qPCR

Total RNA from macrophages treated as described in Figure legends was isolated using Qiagen RNAeasy Plus Mini kit. Total colon RNA was isolated using Trizol (Life Technologies) reagent and gene expression assessed using the standard curve method for relative quantification. RNA samples were subjected to random-primed cDNA synthesis and gene expression was analyzed by qPCR with Maxima Sybr Green/ROX/2x master mix (Thermo Scientific^Tm^) on StepOne Plus real time PCR system (Applied Biosystems) using standard protocols, and data was analyzed using δδCt method with b-actin transcript as housekeeping control. Primers are listed in Extended Data Table 4.

### RNAseq

The Quality of total RNA was assessed using Agilent BioAnalyzer 2100 for RNA integrity. Samples with RIN>8 were used for library preparations. Sequencing libraries were prepared using the NEBNext Ultra II Directional RNA Library Prep Kit for Illumina, following the manufacturer’s instructions. Libraries quality was evaluated using Agilent BioAnalyzer 2100 and the libraries were sequenced by a NovaSeq 6000 system (50 bp, paired-end) at the Genomics Resources Core Facility at Weill Cornell College of Medicine at 15-25 mln reads per sample. All experiments were performed in at least triplicates.

### ATACseq

ATACseq was performed as described ^65, 66^ with minor modifications. Briefly, 10^5^ cells were lysed in ATAC lysis buffer [10 mM Tris-HCl (pH 7.4), 10 mM NaCl, 3 mM MgCl_2_, and IGPAL CA-630] to isolate nuclei. The resultant nuclei were resuspended in the transposase reaction mix (25 μL x TD buffer (Nextera DNA Sample Preparation Kit), 5 μL Illumina Tn5 transposase, and 22.5 μL nuclease-free water and incubated at 37 °C on a thermomixer at 1000 rpm for 30 min. The transposed DNA was purified using a QIAGEN MinElute Purification Kit, then amplified using NEBNext High-Fidelity 2X PCR Master Mix and unique ATAC indexing primer for 10 cycles. Post-amplification, libraries were double size-selected for removal of primer dimers and DNA fragments (>1000 bp). The amplified libraries were quality-controlled by Agilent BioAnalyzer and pooled. 50-bp paired-end sequencing was performed on an Illumina NovaSeq 6000 system at the Genomics Resources Core Facility at Weill Cornell Medicine. All experiments were performed in quadruplicates.

### ChIP-qPCR and ChIPseq

Macrophages were crosslinked with 1% methanol-free formaldehyde (Thermo) for 10 min (H3K27ac) or double-crosslinked with 2 mM disuccinimidyl glutamate (Proteochem, c1104) for 30 min followed by 1% methanol-free formaldehyde for 10 min at RT (GR) ^36^. Nuclear extracts were prepared ^21^ and chromatin DNA was sonicated using Bioruptor Pico (Diagenode, 8 cycles, 30s on/off bursts) to generate DNA fragments ∼200-700 bp. After sonication, 10% of nuclear extract was saved as input and rest incubated with protein A/G agarose (Santa Cruz) for 30 min at 4 °C. Precleared extracts were incubated with 40 μL of 50% protein A/G agarose and 5 of normal rabbit IgG (Santa Cruz Biotech), H3K27ac (Active Motif, 39133), GR (Thermo Fisher Scientific, PA1-511A) at 4 °C overnight. The following day, beads were washed 8x with modified RIPA buffer containing 500 mM LiCl and 2x with TE + 50 mM NaCl buffer. DNA was extracted after reversing crosslinks and purified with Qiagen PCR purification kit. Resulting DNA was analyzed by qPCR (primer sequences are listed in Supplementary Data 1) with 28S as an internal control and occupancies expressed relative to the normal IgG baseline. For ChIPseq, libraries were prepared using NEBNext Ultra II Library Prep Kit for Illumina (NEB), size selected (150-500 bp), and PCR-amplified for 15 cycles. Libraries were sequenced at the Genomics Resources Core Facility at Weill Cornell Medicine using NextSeq500 (75 bp single-end).

### CUT&RUN

CUT&RUN assay was performed using the EpiCypher CUTANA^TM^ ChIC/CUT&RUN kit v2.1 according to manufacturer’s protocol. In brief, 10^6^ cells were harvested for each condition, washed with 1X PBS, and resuspended in wash buffer, followed by incubation at RT for 10 min with activated Concanavalin A (ConA) beads. Cells were then permeabilized and incubated overnight on a nutator at 4°C with 1 μg of normal rabbit IgG (EpiCypher), KLF4 (proteintech, 1180-1-AP), or GRIP1 (Novus Biologicals, NB100-1756) antibody. Cell-bead antibody complex was then washed with cell permeabilization buffer and incubated with pAG-MNase (EpiCypher), followed by addition of 2 mM CaCl_2_ and incubation for 2 h at 4°C. Stop buffer was added to the reaction mixture and DNA fragments were released and purified with a CUTANA DNA purification kit (EpiCypher). DNA library construction was performed using the NEBNext Ultra II DNA Library Prep Kit from NEB (E7645L). The libraries were sequenced on Illumina NovaSeq 6000 system (50 bp paired-end) at the Genomics Resources Core Facility at Weill Cornell Medicine.

### Phagocytosis assay

Macrophages were seeded into a 96-well plate (80,000 cells/well) and polarized with IL4, Cort or Dex for 24 h. After removing the media, 0.75 mg/ml pHrodo *S. aureus* Green BioParticles (ThermoFisher Scientific, P35367) were added for 1 h at 37°C. Before imaging, macrophages were stained with 1 μM Hoescht dye for 5 min at 37°C. The plates were read on the ImageXpress Micro Confocal Imager at the Weill Cornell Microscopy and Image Analysis Core. Fluorescence puncta count and intensity were measured and normalized to the macrophage count for each image. Background fluorescence intensity and puncta count from macrophages alone and BioParticles alone were subtracted.

### Acute DSS colitis histopathology

Mice were given 2 % dextran sulfate sodium (DSS) (MP Biomedicals) in drinking water, evaluated daily for body weight changes, and sacrificed when at least 1 mouse lost 20 % of body weight (2-3 weeks). Colons were harvested and processed for histopathology or total RNA isolation. The colon tissues were fixed in 10 % formalin and histopathological changes of colons were assessed (Tania Pannellini, HSS) using H&E staining (Laboratory of Comparative Pathology, MSKCC) and graded as follows: Grade 1 – well defined crypts and villi; mucus secretion is preserved; mild lymphocytic inflammation; Grade 2 – villi are short and distorted, crypts are progressively lost, goblet cells are diminished, hyperplasia of glandular and superficial epithelium; Grade 3 – crypts are lost, ulceration of mucosa, severe inflammation of mucosa and submucosa, heavy infiltration of lymphocytes, macrophages and granulocytes. Surface area occupied by histopathological changes of each grade was quantified.

### Data analysis RNAseq

Read quality filtering and adapter trimming were performed with fastp 0.23.2 ^67^. Filtered reads were mapped to the mouse genome (mm10) with the STAR aligner 2.7.0 ^68^ and exonic reads were counted against GENCODE release 27 annotation with featureCount ^69^ using a customised pipeline available at https://gitlab.com/hssgenomics/pipelines/RNAseq. Differential gene expression analysis was performed with edgeR 3.38.4 ^70^ using quasi-likelihood framework. Genes with low expression levels (<2 counts per million in at least one group) were filtered from all downstream analyses. P values were adjusted to correct for multiple testing using the Benhamini-Hochberg FDR procedure. Genes with adjusted p-values < 0.05 and log2 fold change >1 were considered differentially expressed. Downstream analyses were performed in R using a Shiny-driven visualization platform (RNAseq DRaMA, 1.4.2, https://gitlab.com/hssgenomics/Shiny) developed at the HSS David Z. Rosensweig Genomics Research Center. Differentially regulated pathways were identified using R QuSAGE 2.30.0 package ^71^ with MSigDB C2 set (curated gene sets c2.cp.v7.3). All gene set with less than 10 genes were not included in the analysis. Pathways with unadjusted p <0.05 were selected for initial analysis and were further filtered with FDR < 0.1 to correct for multiple testing.

### ATACseq

Paired-end reads are pre-processed using fastp which supports adapter trimming, low quality base trimming, and calculation of additional quality control metrics. Trimmed reads were aligned to the mm10 genome using bowtie2 ^72^ with *--very-sensitive-local –q –p* options set. Low quality, improper, multimapping alignments, and PCR duplicates are filtered using SAMtools ^73^. Peaks are called on each sample (replicas are not pooled) with MACS2 ^74^ using *macs2 callpeak –-nomodel –-shift –100 –-extsize 200 –q 0.05* options. After peak calling, all peaks from all samples called separately were combined and adjacent peaks (within 10bp) were merged. Reads from each sample within the combined set of peaks are then counted using featureCounts to produce an initial reads-per-peak count matrix. The ATACseq/ChIPseq pipeline is publicly available at https://gitlab.com/hssgenomics/pipelines/ATACseq.

The quality of individual experiments was evaluated by calculating Fraction of reads in Peaks (FRiP), fragment size distribution, analyzing MACS2 peak statistics distributions (peak width, pileup and enrichment over the background). Furthermore, to ensure consistency read-in-peak pairwise Spearman correlations were determined within experimental groups. The average within group correlation was larger than 0.95 (Extended Data Table 5).

The reads-per-peak counts were analyzed for differential peaks using the edgeR quasi-likelihood framework which allows for building complex modeling solutions not attainable by peak calling algorithms at the peak calling step (*e.g.*, with MACS2 directly). Before the analysis, we removed poorly reproducible peaks containing zero reads in more than 3 samples within a group and peaks with the length of more >1000 base pairs. Modeling the replicated ATACseq data using the counts-per-peak framework allows us to calculate logFC, p-values, and FDR for each peak. All peaks were annotated to the nearest gene and TSS, if the peak overlaps an annotated Fantom5 ^75^ enhancer, that enhancer is reported. Peaks with FDR<0.05 were considered differential.

We used HOMER ^31^ to identify overrepresented transcription factor motifs in peak assemblages. To identify genomic locations of individual binding motif identified by HOMER we used *matchMotifs* function from R *motifmatchr* package and position weight matrices (PWM) from JASPAR 2022 ^76^. The 5’ Tn5 cut site counts within binding motifs of interest and surrounding genomics region (−250/+250) were normalized to the library size and to the local background outside motif-containing peaks using *bamProfile* function R *bamsignals* package ^77^.

### ChIPseq

Single-end reads were preprocessed as described in the ATACseq section. Peaks are called on individual samples without replica pooling using *macs2 callpeak macs2 –-broad –-bw 180 –-keep-dup all –-q 0.05* with IgG input as a control. Following peak calling, peaks were merged and reads and peaks were counted with *featureCount* as described in ATACseq section. Differential peaks and overrepresented binding motifs were identified as described in the ATACseq section.

### CUT&RUN

Samples were processed using an in-house nextflow-based pipeline available at https://github.com/michaelbale/jlabflow. Briefly, reads were trimmed using the Cutadapt wrapper trim-galore and mapped to mm10 with bowtie2 using *--very-sensitive-local* with a maximum insert size of 1000 bp. The initial bam file was then filtered for a minimum MAPQ of 30. Subsequently, PCR duplicates and reads mapping to the mitochondrial chromosome or within regions from a unified forbidden list from ENCODE and ^78, 79^. Finally, only correctly mated mapped reads were retained. Peak calls were made using MACS2 on each individual sample and merged using the IDR framework within sample conditions in an iterative manner then merged across sample conditions for GRIP1 or retaining peaks that reciprocally overlapped each other’s lengths by at least 30% for KLF4. Finally, TMM-normalized signal tracks were visualized using deeptools bamCoverage with a calculated scaleFactor derived from the edgeR package *calcNormFactors* function.

Differential peak analysis was carried out using DESeq2 and all peaks with an FDR ≤ 0.05 were called as differential. Average profile plots within differential regions were generated using deeptools computeMatrix and plotProfile with the TMM-normalized signal track bigwigs. The upset plot was generated using the BedSect web-app with input peak sets as indicated. Global peak-set motif enrichment was done as described with HOMER. The motif heatmaps were generated by calculating deviation z-scores from the chromVAR package with the homer_pwms HOMER motifs from chromVARmotifs. Motifs with significant variability (p_adj_ ≤ 1*10^-6^) were retained for heatmaps.

## Acknowledgements

All sequencing for this study was performed at the Weill Cornell Genomics Resources Core Facility. We are grateful to the Center of Comparative Medicine and Pathology of Memorial Sloan-Kettering Cancer Center as well as the Laboratory of Comparative Pathology at Weill Cornell for histopathology analysis. Tania Pannellini (HSS) assisted in the analysis of H&E staining (Laboratory of Comparative Pathology, MSKCC). This work was supported by the grants to I.R.: NIH R01DK099087, NIH R01AI148129 and The Hospital for Special Surgery David Rosensweig Genomics Center, to S.Z.J.: NIH R01AI148416, and the NIH fellowship F1 HL152706-01A1 to A.W.D.

---

**Extended Data Fig. 1:**
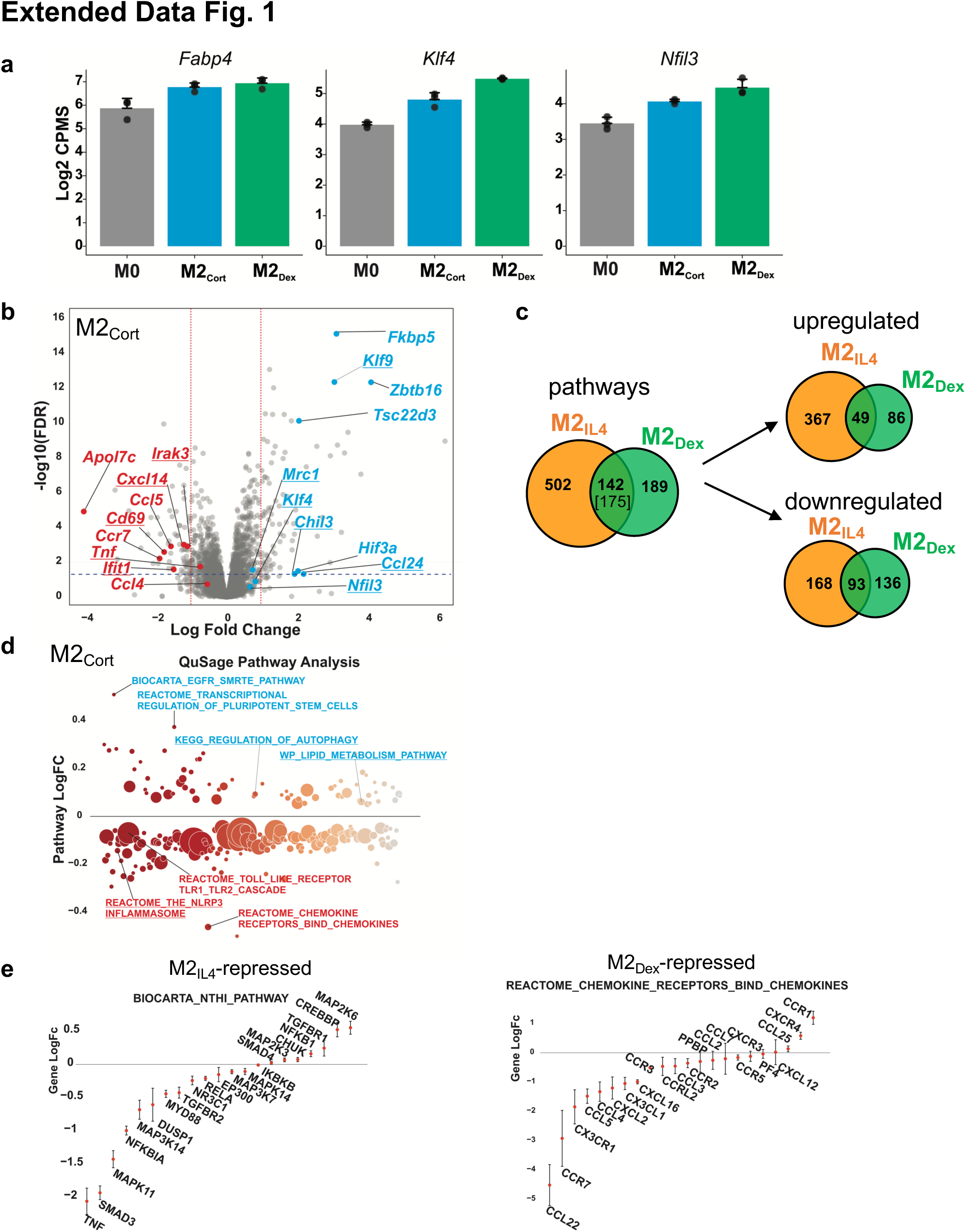
Transcriptomic analysis of the M2_IL4_ and M2_GC_ macrophage populations. **a** Expression of selected genes upregulated in M2_Cort_ and M2_Dex_ (n=3). **b** Volcano plot shows DEGs in M2_Cort_ relative to M0 (LogFC ≥ 1; FDR <0.05). Selected upregulated genes are highlighted blue, and downregulated in red. Some of the shared target genes between M2_IL4_, M2_Dex_, and M2_Cort_ are underlined. **c** Venn diagrams show differentially regulated pathways determined by QuSAGE in M2_IL4_ and M2_Dex_ relative to M0 (p <0.05). **d** Differential pathways in M2 determined by QuSAGE as in Fig. 1d. Underlined are some of the pathways shared between M2_IL4_, M2_Dex_ and M2_Cort_ (upregulated, blue; downregulated, red). **e** Genes from two pathways selectively repressed in M2_IL4_ and M2_Dex_ are plotted as logFC±SD.

**Extended Data Fig. 2:**
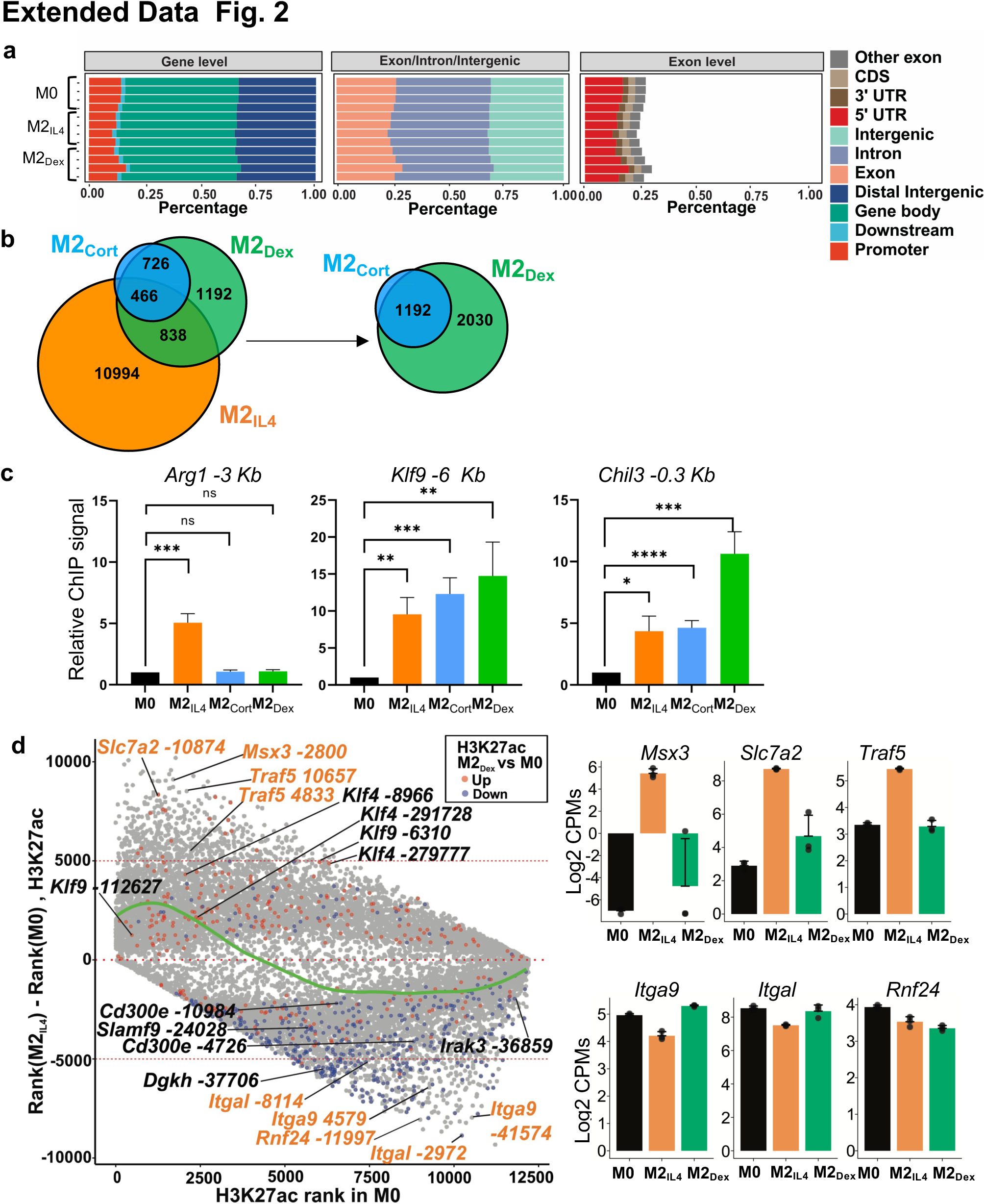
M2_IL4_ and M2_GC_ macrophage populations share epigenetic landscape. **a** The distribution of ATACseq peaks relative to known genomic features in M0, M2_IL4_ and M2_Dex_ populations across replicates (n=3). **b** Venn diagram shows differential M2_IL4_, M2_Cort_ and M2_Dex_ H3K27ac peaks relative to M0. **c** H3K27ac assessed by ChIP-qPCR at enhancers of indicated genes in M2_IL4_, M2_Cort_ and M2_Dex_ normalized to that in M0. Student’s t-test; *p < 0.05, **p < 0.01, ***p < 0.001, ****p < 0.0001, ns, non-significant. n=3, error bars are SEM. **d** Rank shift analysis of mean H3K27ac signals in M0 (baseline) and M2_IL4_. Peaks that are also differentially regulated in M2_Dex_ are colored in red (hyperacetylated) and blue (hypoacetylated). The LOESS regression line (green) shows the relationships between acetylation strength in M0 and signal change in M2_IL4_. Enhancer sites associated with representative IL4-specific (orange) and shared (black) genes are marked, and the expression of representative genes as determined by RNAseq (Fig. 1) is plotted on the right.

**Extended Data Fig. 3:**
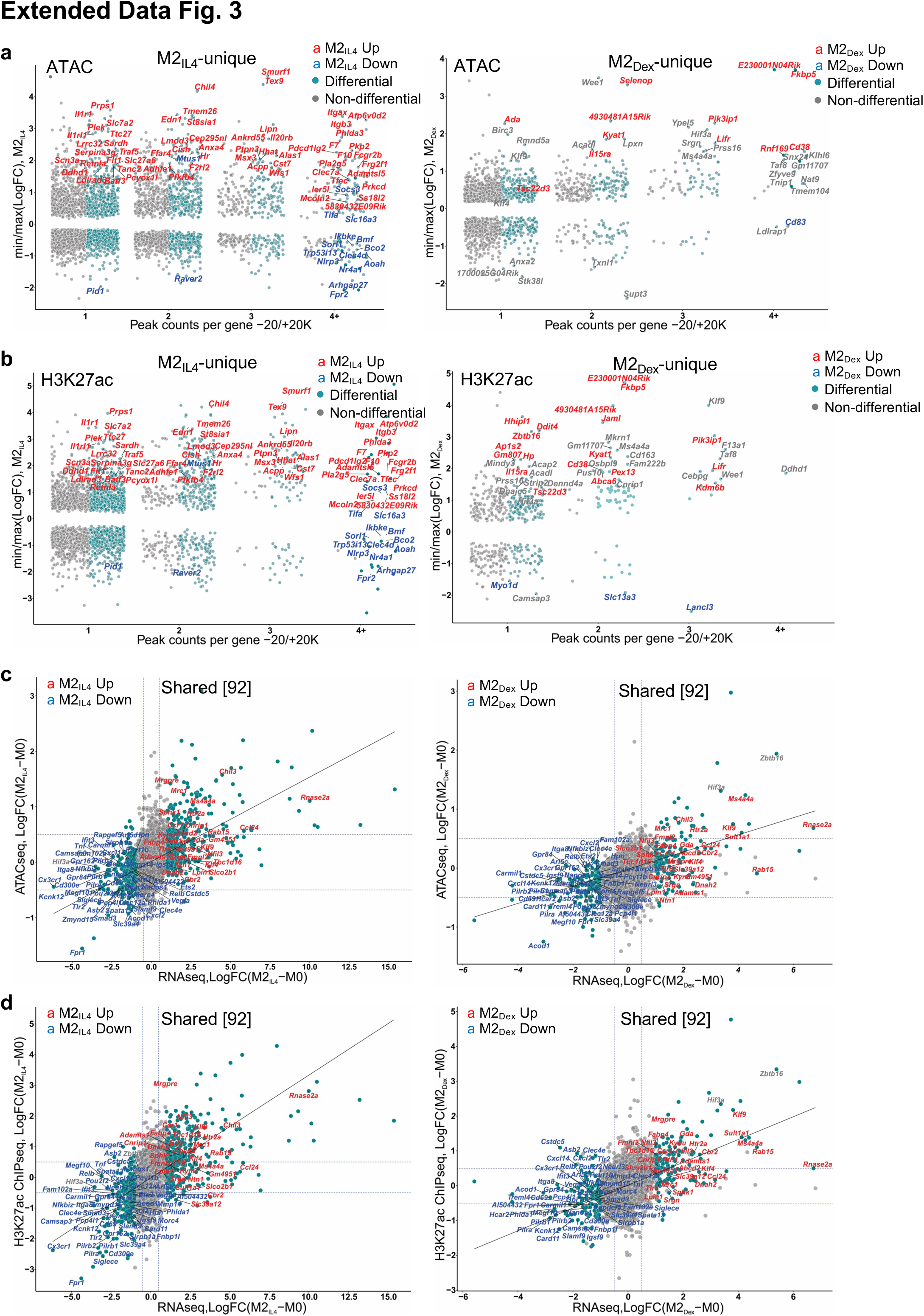
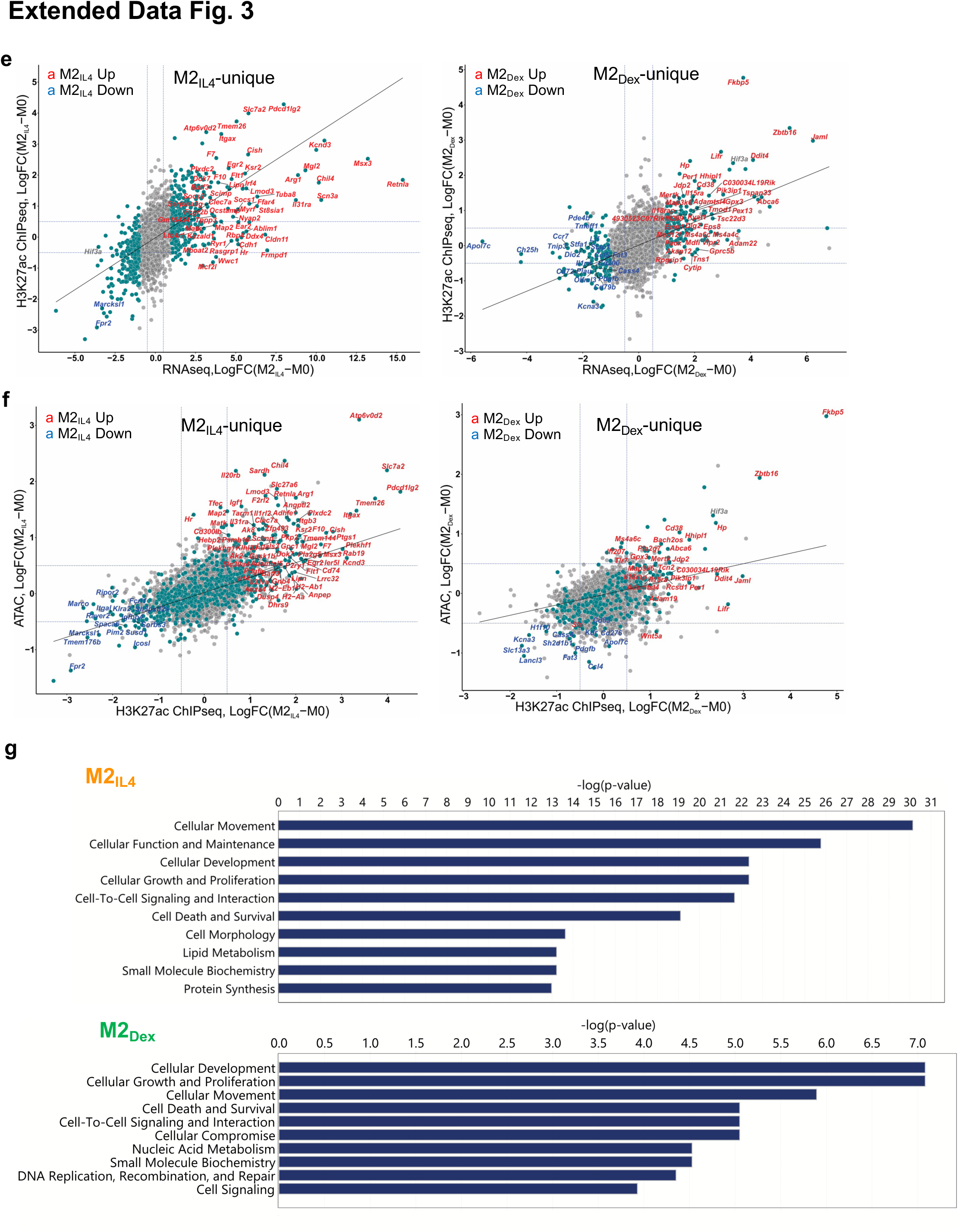
Signal-specific and shared enhancers in M2_IL4_ and M2_GC_ macrophages. **a-b** Comparisons of TSS window-associated ATACseq (**a**) or H3K27ac ChIPseq (**b**) peaks in differential (teal) vs. non-differential (gray) genes in M2_IL4_ (left panels) and M2_Dex_ (right panels) stratified by the number of gene-associated peaks. Only peaks with the largest positive (max) or negative (min) changes relative to M0 are shown. A subset of up-(red) and downregulated (blue) M2_IL4_-unique (left) and M2_Dex_-unique (right) DEGs identified by RNAseq (Fig. 1) are labeled. **c-d** Correlations between normalized RNAseq and ATACseq (**c**) or H3K27ac ChIPseq (**d)** signals in the TSS-proximal – 1K + 1K windows for differential (teal) and non-differential (gray) genes in M2_IL4_ (left) and M2_Dex_ (right). A subset of 92 shared up-(red) and downregulated (blue) DEGs identified by RNAseq (Fig. 1) are labeled. **e-f** Correlations between normalized RNAseq and ATACseq (**e**) or ATACseq and H3K27ac ChIPseq (**f**) signals in the TSS-proximal –1K + 1K windows for differential (teal) and non-differential (gray) genes in M2_IL4_ (left) and M2_Dex_ (right). A subset of the up-(red) and downregulated (blue) 587 M2_IL4_-unique DEGs (left) or 174 M2_Dex_-unique DEGs identified by RNAseq (Fig. 1) are labeled. **g** Differentially regulated pathways identified by Ingenuity Pathway Analysis (QIAGEN) of genes associated with change in both the ATACseq and H3K27ac ChIPseq signal in M2_IL4_ and M2_Dex_ populations.

**Extended Data Fig. 4:**
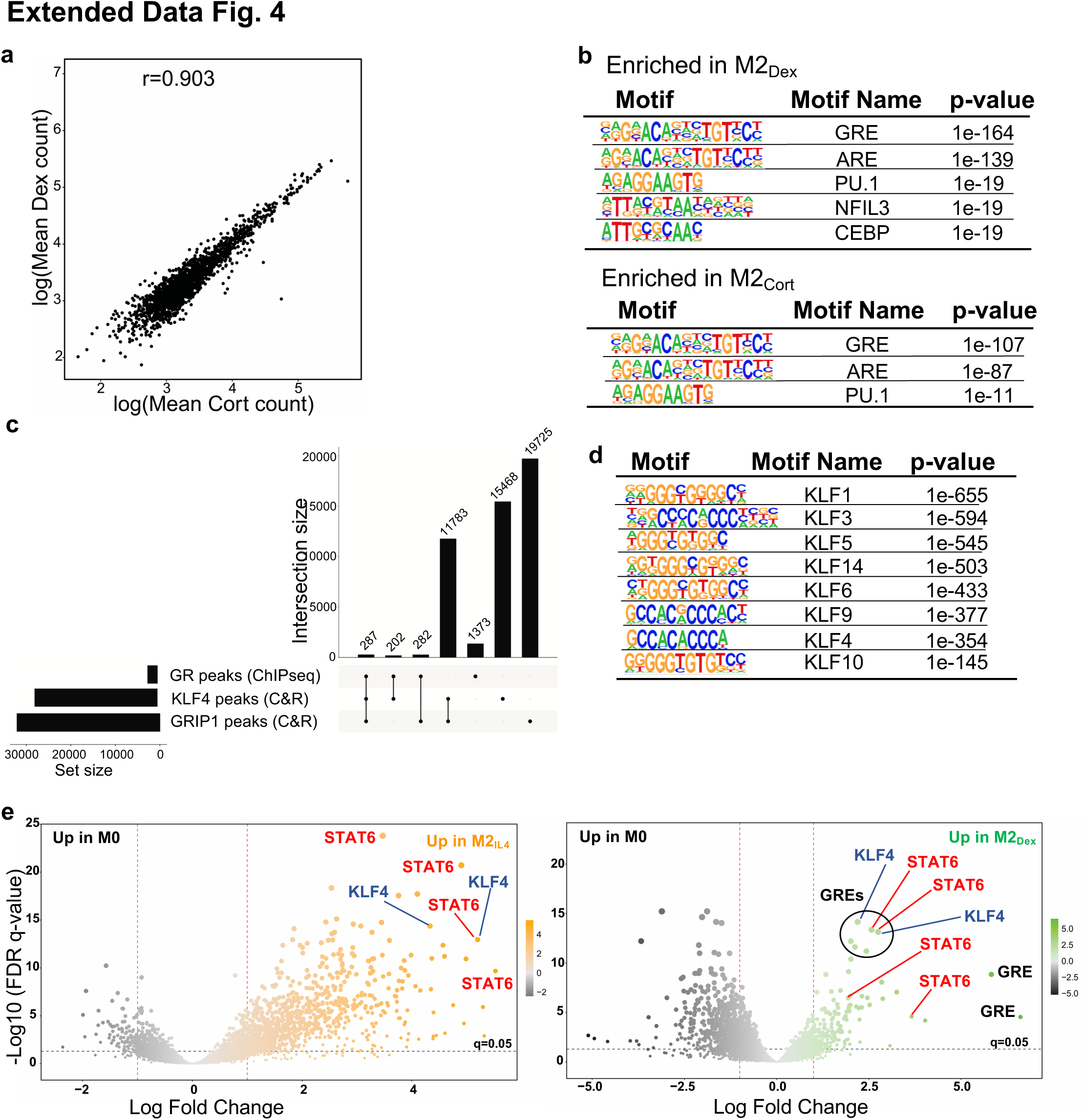
GR, KLF4 and GRIP1 genome-wide binding in differentially polarized macrophage populations. **a** Read counts in peaks Spearman’s correlation between M2_Dex_ and M2_Cort_ GR ChIPseq. **b** Top enriched transcription factor binding motifs in GR ChIPseq peaks in M2_Dex_ and M2_Cort_. Peaks were analyzed by HOMER2 findMotifsGenome.pl to identify significantly enriched motifs relative to genomic background. **c** UpSet plot shows GC-induced peaks from GR ChIPseq and the total peak atlases from KLF4 and GRIP1 CUT&RUN as analyzed by *BedSect*. Set-size: number of total regions (overlaps and unique). Intersection size: number of total regions with specified overlaps. **d** Enrichment p-values for KLF family motifs in KLF4 CUT&RUN-derived peaks. Peaks were analyzed by HOMER2 findMotifsGenome.pl to identify significantly enriched motifs relative to a set of background regions with similar GC content. **e** Volcano plots of GRIP1 CUT&RUN differential peaks in M2_IL4_ (left) and M2_Dex_ (right) relative to M0. Size of the peak point on graph is commensurate with – log10(FDR), whereas color shade is proportional to the log2FC. Peaks were scanned for the presence of transcription factor binding motifs within the HOMER database using *motifmatchr* and select motifs (GRE, black; STAT6, red; KLF4, blue) are indicated.

**Extended Data Fig. 5:**
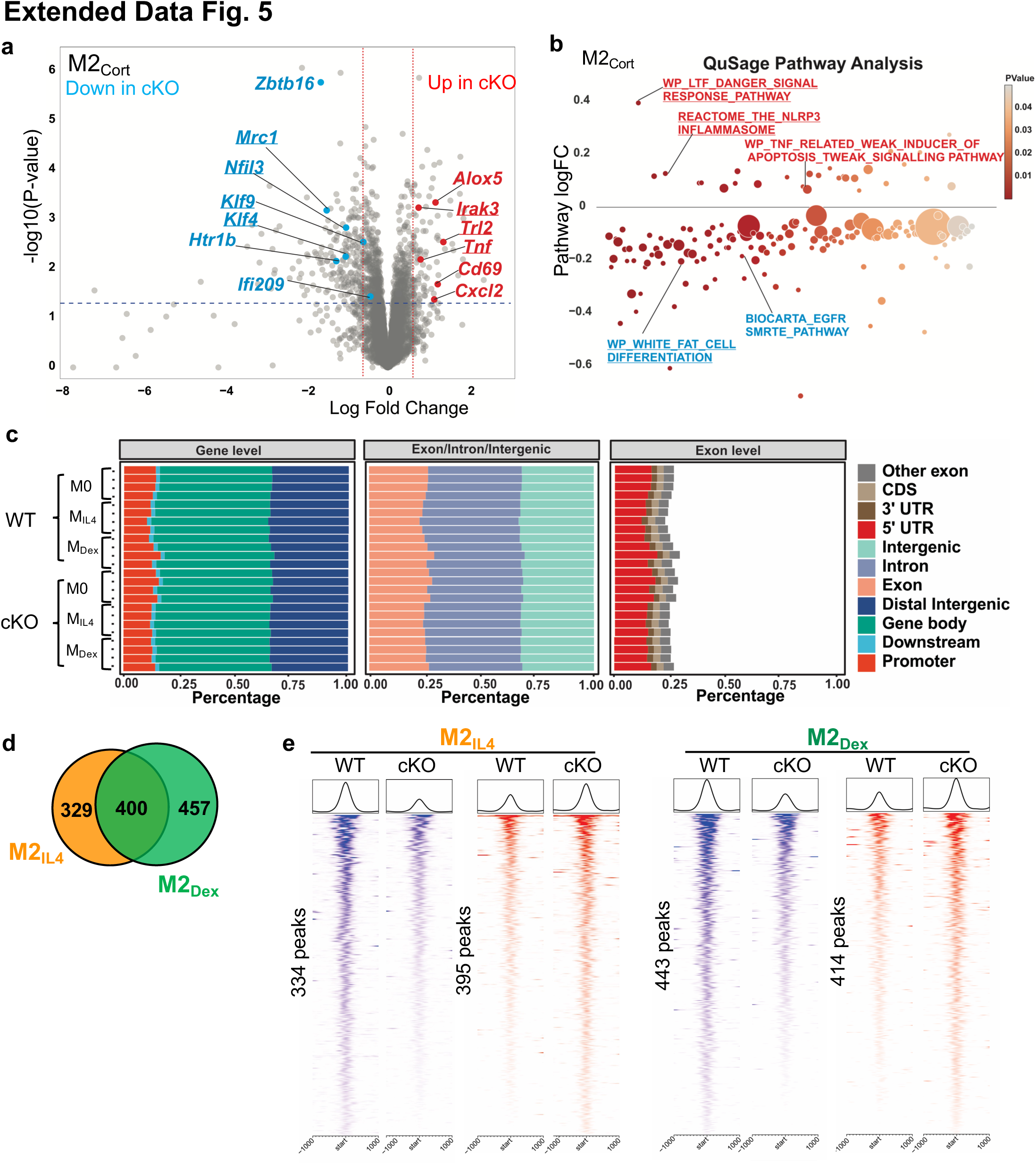
GRIP1 is required for the M2_IL4_ and M2_GC_ programming. **a** Volcano plot shows DEGs in GRIP1 cKO vs. WT in M2_Cort_ relative to M0 of each genotype. (FC ≥1.5; unadjusted p-value<0.05). A subset of DEGs expressed at higher levels in the cKO relative to WT are highlighted in red, whereas those downregulated – in blue. Examples of the shared M2_IL4_:M2_GC_ GRIP1 target genes are underlined. **b** Differentially regulated pathways in M2_Cort_ (unadjusted p < 0.01) identified by QuSAGE with MsigDB are shown for GRIP1 cKO vs. WT. Select up-(red) and downregulated (blue) pathways are labeled, and those shared between IL4 and GC are underlined. Circle size is proportional to the number of genes in the pathway and color signifies p-value. **c** The distribution of ATACseq peaks relative to known genomic features in M0, M2_IL4_ and M2_Dex_ populations across replicates for WT (n=3) and GRIP1 cKO (n=4). **d** Venn diagram of GRIP1-dependent differential ATACseq peaks in M2_IL4_ and M2_Dex_ (n=4, FC ≥ 2; FDR <0.05). **e** Heatmaps of the GRIP1-dependent ATACseq peaks in M2_IL4_ and M2_Dex_ (FC ≥ 2; FDR <0.05) are separated into those dependent on GRIP1 for opening (blue) or closing (red).

**Extended Data Fig. 6:**
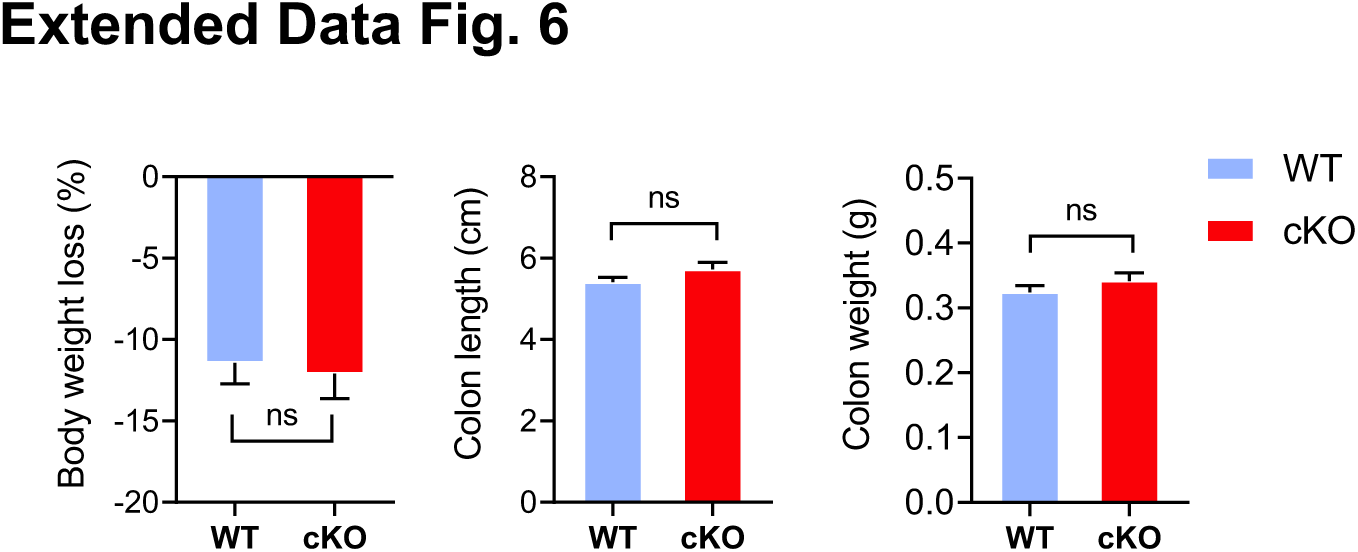
GRIP1 contributes to phagocytic activity of M2_IL4_ and M2_GC_ *in vitro* and healing properties *in vivo*. Body weight, colon length and weight changes of WT and cKO mice 2 wks after initiating DSS-induced colitis.

